# Sparse Multitask group Lasso for Genome-Wide Association Studies

**DOI:** 10.1101/2024.12.20.629593

**Authors:** Asma Nouira, Chloé-Agathe Azencott

## Abstract

A critical hurdle in Genome-Wide Association Studies (GWAS) involves population stratification, wherein differences in allele frequencies among subpopulations within samples are influenced by distinct ancestry. This stratification implies that risk variants may be distinct across populations with different allele frequencies. This study introduces Sparse Multitask Group Lasso (SMuGLasso) to tackle this challenge. SMuGLasso is based on MuGLasso, which formulates this problem using a multitask group lasso framework in which tasks are subpopulations, and groups are population-specific Linkage-Disequilibrium (LD)-groups of strongly correlated Single Nucleotide Polymorphisms (SNPs). The novelty in SMuGLasso is the incorporation of an additional 𝓁_1_-norm regularization for the selection of population-specific genetic variants. As MuGLasso, SMuGLasso uses a stability selection procedure to improve robustness and gap-safe screening rules for computational efficiency.

We evaluate MuGLasso and SMuGLasso on simulated data sets as well as on a case-control breast cancer data set and a quantitative GWAS in *Arabidopsis thaliana*. We show that SMuGLasso is well suited to addressing linkage disequilibrium and population stratification in GWAS data, and show the superiority of SMuGLasso over MuGLasso in identifying population-specific SNPs. On real data, we confirm the relevance of the identified loci through pathway and network analysis, and observe that the findings of SMuGLasso are more consistent with the literature than those of MuGLasso. All in all, SMuGLasso is a promising tool for analyzing GWAS data and furthering our understanding of population-specific biological mechanisms.

**Author summary:** Genome-Wide Association Studies (GWAS) scan thousands of genomes to identify loci associated with a complex trait. However, population stratification, which is the presence in the data of multiple subpopulations with differing allele frequencies, can lead to false associations or mask true population-specific associations. We recently proposed MuGLasso, a new computational method to address this issue. However, MuGLasso relied on an ad-hoc post-processing of the results to identify population-specific associations. Here, we present SMuGLasso, which directly identifies both global and population-specific associations.

We evaluate both MuGLasso and SMuGLasso on several datasets, including both case-control (such as breast cancer vs. controls) and quantitative (for example, plant flowering time) traits, and show on simulations that SMuGLasso is better suited than MuGLasso for the identification of population-specific associations. In addition, SMuGLasso’s findings on real case studies are more consistant with the literature than that of MuGLasso, which is possibly due to false discoveries of MuGLasso. These results show that SMuGLasso could be applied to other complex traits to better elucidate the underlying biological mechanisms.

## Introduction

Feature selection methods have emerged as a popular way of framing Genome-Wide Association Studies (GWAS) to uncover the genetic underpinnings of complex diseases, such as cancer. GWAS aim at establishing associations between genetic variants, more specifically Single Nucleotide Polymorphisms (SNPs), and the presence/absence of a disease or a quantitative trait [1–3]. However, their ability to identify relevant variants is limited by several difficulties, including the curse of dimensionality, population stratification, linkage disequilibrium, and the lack of stability of feature selection procedures with respect to small changes in the input samples. Consequently, the application of feature selection requires careful attention to mitigate false discoveries. The challenge in this context is optimizing the stability of selection to identify regions of interest while minimizing false positives [4].

Contrary to the assumption in many existing feature selection methods that SNPs associated with a phenotype are shared across diverse populations, numerous studies highlight population-specific genetic associations with certain diseases [5]. Notably, diseases can manifest distinct prevalence patterns across populations, leading to variations in risk variants from one genetic ancestry to another [6]. For instance, multiple studies underscore that Africans and Europeans exhibit dissimilar genes associated with the lactase-persistence phenotype [7], emphasizing the population-specific nature of genetic influences. Moreover, recent research has revealed significant differences in genetic risk factors of type 2 diabetes among East Asian and European individuals, highlighting the importance of considering population-specific genetic architectures in disease studies [8].

In previous work, we have introduced the Multitask Group Lasso (MuGLasso) framework, designating groups as blocks of SNPs in strong Linkage Disequilibrium (LD) and tasks as subpopulations [9]. We demonstrated its effectiveness in stably identifying SNPs associated with breast cancer. Despite its effectiveness, the original MuGLasso design required additional post-processing steps to discern task-specific LD-groups [9], prompting the introduction of a second regularization term to enhance population-specific sparsity.

Hence this paper introduces the Sparse Multitask Group Lasso (SMuGLasso), an extension of MuGLasso aimed at refining the population-specific selection of LD-groups. By combining the 𝓁_1,2_-norm penalty of MuGLasso with an additional 𝓁_1_-norm at the LD-group level, SMuGLasso seeks to improve the precision of LD-groups selection.

We evaluate the performance of SMuGLasso against MuGLasso using simulated data and the DRIVE breast cancer dataset. In addition to these qualitative phenoytpes, we assess both MuGLasso’s and SMuGLasso’s effectiveness on a quantitative *Arabidopsis thaliana* phenotype, further validating our approaches on non-human data with a large number of subpopulations.

Finally, we use enrichment analyzes and protein-protein interaction networks to analyze SMuGLasso’s findings on the DRIVE breast cancer data sets, shedding light on the molecular mechanisms underlying breast cancer tumor growth.

Finally, we compare the stability of SMuGLasso, MuGLasso, and other existing methods in identifying LD-groups and SNPs associated with a phenotype, aiming to provide a comprehensive understanding of the proposed framework’s advantages in addressing the challenges inherent to GWAS.

## Materials and methods

We introduce SMuGLasso, a four-step framework designed to enhance the precision of population-specific causal variants selection. The steps are similar to those of MuGLasso and are outlined as follows:

1. **Populations Assignment**: Each sample is assigned to a genetic population using PCA and k-means clustering. This results in the assignment of each population to an input task within the multitask framework, facilitating a tailored analysis for distinct subpopulations.
2. **LD-Groups Formation**: LD-groups consisting of strongly correlated SNPs are formed using adjclust [10] to alleviate the curse of dimensionality by conducting feature selection at the group level.
3. **Model Fitting with Dual Penalty**: The model is fitted with a regularization comprising two penalty terms. Firstly, the MuGLasso penalty involves an 𝓁_1,2_-norm, fostering sparsity at the LD-group level across all tasks and populations. Secondly, we add in SMuGLasso an 𝓁_1_-norm penalty to enforce sparsity, specifically at the LD-group level for individual populations. To address computational complexity, the optimization problem is solved using coordinate descent with gap safe screening rules [11].
4. **Stability Selection**: To improve the robustness of the algorithm, we incorporate a stability selection procedure [12] to ensure a more stable genetic variants selection, contributing to the overall resilience of SMuGLasso.

Unlike MuGLasso, the proposed setting eliminates the need for additional post-processing steps to obtain population-specific LD-groups. SMuGLasso stands out by offering a more precise and refined approach to the selection of population-specific causal SNPs, thereby streamlining the process and enhancing the accuracy of the analysis.

As the population assignment, LD-groups formation, and stability selection procedures are similar to those presented in MuGLasso, we refer the reader to [9] for details and proceed with a detailed discussion of the SMuGLasso model fitting itself.

### Notations

Given a set of *p* SNPs measured on *n* samples, we split the *n* samples in *T* subpopulations/tasks, each of size *n*_*t*_ for *t* = 1, …, *T*, and the *p* SNPs in *G* LD-groups, each of size *p*_*g*_ for *g* = 1, …, *G*. For each population *t*, we denote by 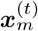 the *p*-dimensional vector of SNPs of the *m*-th sample in the population (*m* = 1, …, *n*_*t*_), and by 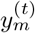 its phenotype.

### MuGLasso and its formulation

In what follows, we recall the formulation of MuGLasso as presented in [9]. MuGLasso leverages a penalized regression framework to model the relationship between SNPs and phenotypes. The formulation seeks to achieve sparsity at the LD-group level and smoothness of regression coefficients within and across tasks. We formulate MuGLasso optimization problem as follows:

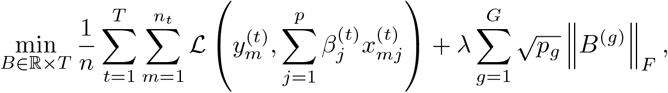

where ***β***^(*t*)^ ∈ ℝ^*p*^ represents the regression coefficients specific to task *t*, denoted as ***β***^(*t*)^ = (*B*_1*t*_, …, *B*_*pt*_). The loss function L takes the form of quadratic loss for quantitative phenotypes (*y* ∈ ℝ) and logistic loss for qualitative phenotypes (*y* ∈ 0, 1). The Frobenius norm ∥·∥*_F_* is used to quantify the size of matrices, and *B*^(*g*)^ is a *p*_*g*_ × *T* matrix containing the regression coefficients for the SNPs of group *g* across all tasks.

MuGLasso can be reformulated by transforming the original dataset into a new one, represented as a block-diagonal matrix denoted as 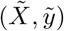. Here,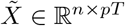 forms a block-diagonal matrix where each of the *T* diagonal blocks corresponds to the SNP matrix 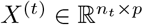 for task *t*. Additionally, 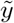 is an *n*-dimensional vector obtained by stacking the phenotype vectors for each task. Introducing this transformation, we derive an adjusted optimization problem. Let 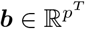 be the vector of regression coefficients, where *p*^*T*^ represents the total number of features across all tasks. The reformulated optimization problem is then:

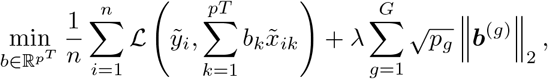

Where 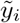 is the *i*-th entry of the transformed phenotype vector 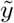, and 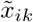 is the (*i, k*)-th entry of the block-diagonal matrix 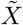.Additionally, 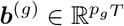 denotes the regression coefficients associated with SNPs of group *g*, and 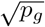 is the square root of the size of group *g*.

### SMuGLasso

#### Problem forumation

To address potential limitations of the MuGLasso framework, we introduce the SMuGLasso method, which incorporates an additional 𝓁_1_ penalty. This penalty aims to improve the selection procedure of specific population LD-groups and discard false positive discoveries, enhancing the identification of relevant loci. The details of the model and its implementation are provided in S1 Appendix. The optimization problem of SMuGLasso is written as follows:

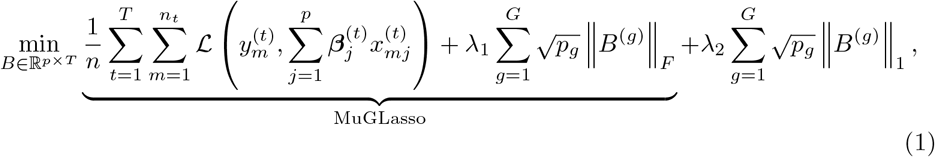

The penalization parameters *λ*_1_ and *λ*_2_ control the respective strengths of the two regularization terms.

#### Optimization

The formulation of SMuGLasso can be transformed exactly as that of MuGLasso shown above. The reformulated model is then:

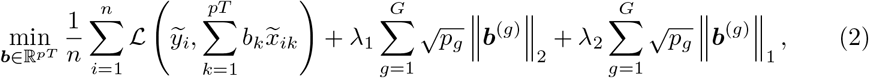

where 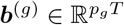 is the vector of regression coefficients corresponding to all SNPs of group *g* for all tasks.

#### Gap safe screening rules

Gap safe screening [11] is a method designed to enhance the efficiency of solving regularization problems in statistical learning and high-dimensional data analysis. Hence, gap-safe employs a set of rules to identify and eliminate irrelevant features from the optimization problem, significantly reducing computational complexity. These rules use duality gaps to rigorously guarantee that the discarded features have zero coefficients in the optimal solution, ensuring the accuracy of the model while improving computational speed. This approach is particularly useful in cases with large datasets and numerous features, making it a valuable tool in GWAS data analysis. We have detailed the fundamentals of these rules in MuGLasso paper [9]. Code is available in https://github.com/asmanouira/SMuGLasso

### Related work

Our method is related to the group Lasso and multitask Lasso, which both rely on an 𝓁_1,2_-norm penalty [13, 14]. Building on that, several studies have been proposed related to multitask variants composed of either two or three regularization terms [15–18].

Notably, these models exhibit limitations in scalability when confronted with high-dimensional data, rendering them inapplicable to our specific context.

To effectively select the additional population-specific regularization term for SMuGLasso, we conducted a comprehensive investigation into the applicability of existing methods. Notably, we found that the proposed sparsity-enforcing penalties were not suited to our specific problem. Our objective is to implement a regularization term that enforces sparsity for specific populations at the level of LD-groups.

We specifically examine the method proposed by (Li L, et al.) [18], which suggests implementing three regularization-based multitask models. Their optimization problem is reformulated as follows:

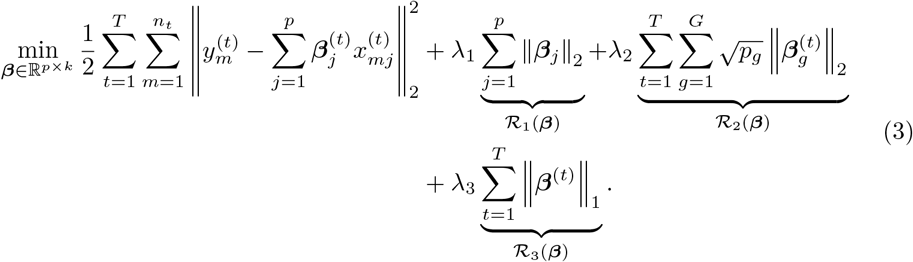

Here, the authors aim to enforce population-specific group sparsity using the term ℛ_2_(***β***), aiming to select certain groups only for specific tasks or subpopulations.

However, relying only on this regularization term results in the optimization problem being separated across tasks, meaning that the selection is performed independently for each single task. To ensure simultaneous task fitting, the authors introduce the term ℛ_1_(***β***), corresponding to multitask regularization at the single-SNP level across all *T* tasks. Additionally, they seek to enforce sparsity within groups using an 𝓁_1_-norm over all SNPs, represented by a third regularization term (defined by ℛ_3_(***β***)), corresponding to the second regularization term of the well known sparse group Lasso.

In our study, we aim to enhance the selection for population-specific LD-groups. Hence, as mentioned above integrating ℛ_2_(***β***) into SMuGLasso would not maintain multitasking across the tasks *T*, while implementing ℛ_3_(***β***) alongside SMuGLasso would impede the interpretability of the selected features. Selecting SNPs within groups for specific populations complicated determining the number of SNPs within an LD-group *g* that must be set to 0 to consider the group as not selected for a particular task *t*.

Additionally, the inclusion of two penalties in SMuGLasso substantially increased computational demands at a GWAS scale.

Another approach has also been proposed presenting a sparse group multitask feature selection model for GWAS data aimed at leveraging pleiotropy, i.e., SNPs associated with multiple complex diseases [19]. However, it’s important to note that their method addresses a different scenario from ours. Specifically, they focus on scenarios where the tasks are output phenotypes rather than input populations samples. In their setting, the goal is to select groups of SNPs targeting the same gene or pathway. Additionally, they combine multiple GWAS datasets and retain only the SNPs shared between them, resulting in a substantial reduction in the number of SNPs (down to 3, 766 SNPs) and thereby reducing computational complexity significantly.

## Experiments

### Data

#### Simulated data

Using GWAsimulator [20], we simulate GWAS data following LD patterns of two populations (CEU: Utah residents with Northern and Western European ancestry and YRI: Yoruba in Ibadan, Nigeria) from HapMap3 [21]. We generate different numbers of samples through subpopulations to mimic the structure of real data, where samples through subpopulations are not necessarily equally distributed. We also produce the population stratification confounder by varying the case:control ratio within each subpopulation (CEU 1 300:1 700 and YRI 400:600). We predefine a total of 200 disease SNPs as shown in Table 1, in which 50 SNPs (respectively 50 SNPs) are specific to the CEU (respectively YRI). We locate the predefined disease loci (and their corresponding LD-groups) randomly and without loss of generality through chromosomes 12, 19, 21 and 22, as shown in Table 2. In total, the data is composed of 4 000 samples and 50 000 SNPs. For CEU, there are 1,407 LD-groups, each containing an average of 35 SNPs. For YRI, there are 995 LD-groups, each containing an average of 50 SNPs.

**Table 1.** For simulated data, number of predefined causal SNPs.

| Populations | Number of SNPs |
| --- | --- |
| Specific-CEU | 50 |
| Specific-YRI | 50 |
| Shared (CEU+YRI) | 100 |
| Total | 200 |

**Table 2.** For simulated data, location of predefined disease loci represented by start/end positions and its corresponding LD-groups number in each subpopulation through chromosomes: 12, 19, 21 and 22.

| Chromosome | Subpopulations |  |
| --- | --- | --- |
|  | CEU<br>loci (# LD-groups) | YRI<br>loci (# LD-groups) |
| 12 | 4 000 - 4 050 (3) | 4 000 - 4 050 (1) |
| 19 | 1 000 - 1 050 (2) | 1 000 - 1 050 (2) |
| 21 | ∅ | 10 000 - 10 050 (1) |
| 22 | 1 000 - 1 050 (3) | ∅ |

#### DRIVE Breast Cancer OncoArray

The DRIVE OncoArray dataset contains 28 281 individuals that were genotyped for 582 620 SNPs. 13 846 samples are cases and 14 435 are controls. The dataset contains data for the following countries: USA, Uganda, Nigeria, Cameroon, Australia and Denmark. Additional information about data access and ethical approval are presented in S2 Appendix.

#### Arabidopsis thaliana

Our *Arabidopsis thaliana* dataset comes from the 1001 Genomes Project [22] (Build TAIR10). We study the DTF3 phenotype, which is the time until the first open flower, in days. The dataset obtained from easyGWAS [23] contains 923 samples and 6 973 565 SNPs divided into 5 chromosomes. This dataset contains plant samples coming from 44 countries.

### Preprocessing

#### Quality control and imputation

For the simulated dataset and DRIVE breast cancer, we exclude SNPs with a minor allele frequency below 5%, a p-value for Hardy-Weinberg Equilibrium in controls below 10^−4^, or a genotyping rate missing more than 10%. We also remove duplicate SNPs, as well as samples with over 10% missing SNPs. We impute missing genotypes in DRIVE using IMPUTE2 [24].

For *Arabidopsis thaliana*, we perform the quality control steps recommended by [23]. We use a Box-Cox transformation [25] of the phenotype to improve the measurements normality. We remove SNPs with a minor allele frequency lower than 5%.

#### LD pruning

We perform LD pruning using PLINK [26] with an LD cutoff of *r*^2^ *>* 0.85 and a sliding window of 50Mb for the simulated data and DRIVE. For *Arabidopsis thaliana*, we use an LD cutoff of *r*^2^ *>* 0.75 and a window size of 50Mb. After preprocessing steps, 50 000 SNPs remain in the simulated data, 312 237 SNPs in DRIVE and 564 291 SNPs in the *Arabidopsis thaliana* data.

#### Population structure

We use PLINK [26] to compute the principal components of the genotype matrix. In the simulated dataset, we find two populations, corresponding to the CEU and YRI populations (see Subfigure 9a). In DRIVE, we identify two populations (see Subfigure 9b) that we call in this paper POP1 (samples from the USA, Australia and Denmark) and POP2 (samples from the USA, Cameroon, Nigeria and Uganda).

In the *Arabidopsis thaliana* dataset, among 44 countries, we retrieve 5 populations using k-means clustering of the top 4 principal components (see S4 Fig and S5 Fig). These 5 populations are detailed in S6 Table.

#### LD-groups choice

For simulated and DRIVE data, we determine the LD-groups for each subpopulation and each chromosome using adjclust [10]. However, for *Arabidopsis thaliana*, adjclust did not scale computationally to the huge number of SNPs in the five chromosomes. Thus, we first split each chromosome into independent LD-blocks using snpldsplit [27] function from bigsnpr R package [28]. We then form the LD-groups by applying adjclust on the obtained chunks of independent LD-blocks.

For all three datasets, we then combine these LD-groups across populations by merging their boundary coordinates to obtain shared LD-groups. Table 3 shows the number of LD-groups obtained for each subpopulation and the final number of shared groups.

**Table 3.** Number of LD groups for each subpopulation of the studied datasets (simulated, DRIVE and *Arapidopsis thaliana*), and after combination across subpopulations.

| Data | Subpopulations | # LD-groups | # shared LD-groups |
| --- | --- | --- | --- |
| Simulated data | CEU | 1 407 | 1 566 |
|  | YRI | 995 |  |
| DRIVE real data | POP1 | 8 152 | 17 782 |
|  | POP2 | 5 032 |  |
| *A. thaliana* data | POP1 | 1 846 | 7 080 |
|  | POP2 | 1 950 |  |
|  | POP3 | 2 002 |  |
|  | POP4 | 1 728 |  |
|  | POP5 | 1 834 |  |

### Comparison partners

As a baseline, we use PLINK to conduct association studies between each SNP individually and the phenotype, employing either the top PCs as covariates (**Adjusted GWAS**) or treating each population separately (**Stratified GWAS**). Additionally, we derive a PCA-adjusted phenotype by regressing the top PCs against the phenotype to compute the residuals. To explore the impact of grouping correlated SNPs, segregating populations into tasks, and using an additional penalty to automatically select population-specific SNPs, we compare SMuGLasso with MuGLasso and various other methods. These include a single-task Lasso without groups applied to each population separately (**Stratified Lasso**) or the adjusted phenotype (**Adjusted Lasso**), as well as a single-task group Lasso **Stratified group Lasso** and **Adjusted group Lasso** applied similarly with an adjusted phenotype.

Note that for **Adjusted Lasso** and **Adjusted Group Lasso**, when handling qualitative phenotypes (such as case-control in DRIVE), we employ logistic regression on the top principal components (PCs) for adjustment. The resulting residuals are subtracted from the actual phenotype values (1 for cases or 0 for controls) to generate a newly adjusted phenotype. Similarly, for quantitative phenotypes (e.g., DTF3 in *Arabidopsis thaliana*), we apply the same procedure but with linear regression. Traditionally, PCA-based methods or linear mixed models are commonly used for population stratification adjustment in GWAS. For instance, FastLMM [29] is recommended for *Arabidopsis thaliana*. However, integrating linear mixed models for feature selection poses challenges in machine learning applications. With such approaches, it is always possible to add top PCs as additional features, but there is no guarantee that the method will select, and therefore use, them.

In practice, we use bigLasso [30] for the lassos and gap safe screening rules [11] for the group Lasso to optimize computational efficiency. Across all methods, we determine the regularization hyperparameter(s) through cross-validation. More specifically, we use f1-score as a criterion for binary phenotypes and RMSE for quantitative phenotypes.

To assess methodological performance, we analyze runtime, the ability to identify true causal SNPs (in simulated data), and the stability of feature selection. To quantify the stability of the feature selection procedure with respect to perturbations of the input, we repeat the feature selection process on 10 subsamples of the data and report the average Pearson’s correlation among all pairs of indicator vectors representing the selected features for each subsample (see S1 Appendix).

### Biological interpretation

#### Functional mapping and annotations analysis

We use FUMA [31] to map functionally annotated SNPs to genes according to the physical position in the genome and eQTL mapping. The tool uses information from multiple biological data to perform these mapping analyses. We used Ensembl version 110 as the reference, and the 1000 Genome Project/Phase3 as the reference panel. A physical mapping window of 10 kb was employed to map variants to nearby genes. We set the significance threshold at a p-value of 5.10^−8^ and the eQTL mapping was performed using GTEx data version 6, including all available tissue types.

For *Arabidopsis thaliana* dataset, we map SNPs identified by Adjusted GWAS, MuGLasso, and SMuGLasso to genes using TAIR10, which provides genomic location data of *Arabidopsis thaliana* genes in GFF3 format.

#### Gene set enrichment analysis

We use Metascape [32] to perform gene set enrichment analysis to understand the functional relevance of the obtained gene lists that Adjusted GWAS, MuGLasso and SMuGLasso have discovered. This tool performs pathway and process enrichment analysis using multiple pathway data bases: KEGG Pathway, Reactome Pathway, WikiPathways, PID, BioCarta, Panther Pathway, SMPDB, GO Biological Processes, CORUM, TargetScan Pathway, TF targets and PharmGKB. Metascape also performs gene set enrichment analysis against cell type signatures, the gene-disease database DisGeNET, the pattern gene database PaGenBase, and transcription factor targets. Metascape collects terms with an enrichment p-value *<* 0.01, a minimum number of occurrences of 3, and an enrichment factor *>* 1.5 and groups them into clusters based on membership similarities. The term with the smallest p-value within a cluster then represents the cluster. For *Arabidopsis thaliana*, we also use Metascape with the same parameters by specifying A.thaliana as a species.

#### Protein-protein interaction network analysis

We further use Metascape [32] to construct a protein-protein interaction network between enriched genes. Metascape uses multiple sources to this end, including experimental and predicted interactions, and identifies densely connected network components using MCODE [33]. We finally visualize the obtained network using Cytoscape [34] to enable the discovery of functionally related gene groups within breast cancer disease on the DRIVE dataset and on within DTF3 on the *Arabidopsis thaliana* dataset.

## Results

### SMuGLasso and MuGLasso rely on both LD-groups and the multitask approach to recover disease SNPs

On simulated data, we observe that SMuGLasso and MuGLasso outperform the other methods at recovering the predefined disease loci (See Figure 1). In addition, we confirm that performing feature selection at the level of LD-groups provides better performance compared to the conventional single-SNP selection. This confirms that grouping SNPs helps to alleviate the curse of dimensionality and improve the identification of causal variants.

**Fig 1.**
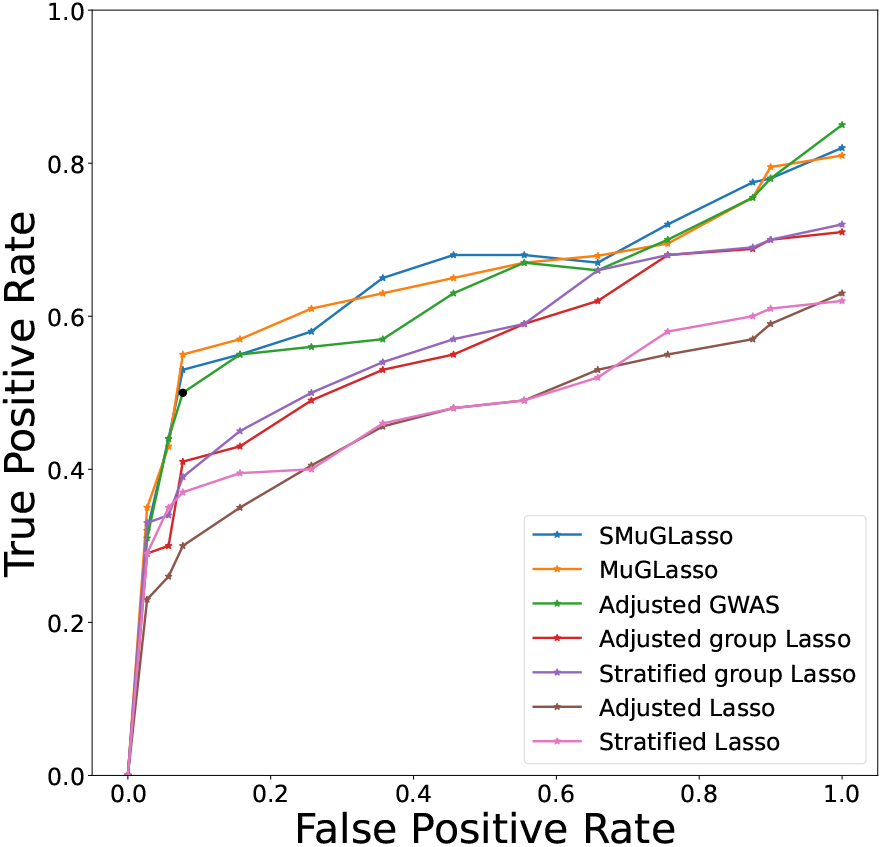
On simulated data, ability of different methods to retrieve causal disease SNPs as a ROC plot

Table 6 gives, for both SMuGLasso and MuGLasso, the number of selected LD-groups and SNPs across and per subpopulation for each dataset. Compared to MuGLasso, we notice that SMuGLasso ensures more sparsity for shared selection across all subpopulations thanks to its additional 𝓁_1_-norm penalty.

SMuGLasso provides a more precise selection for population-specific level. Indeed, SMuGLasso recovers successfully causal LD-groups/SNPs that MuGLasso missed in simulated data (see Figure 5).

We note that SMuGLasso is more intensive computationally compared to MuGLasso and any other tested method (see Figure 2). This computational cost is caused by the additional populations-specific regularization term. However, the implementation is efficient enough to scale to high-dimensional GWAS data thanks to gap-safe screening rules.

**Fig 2.**
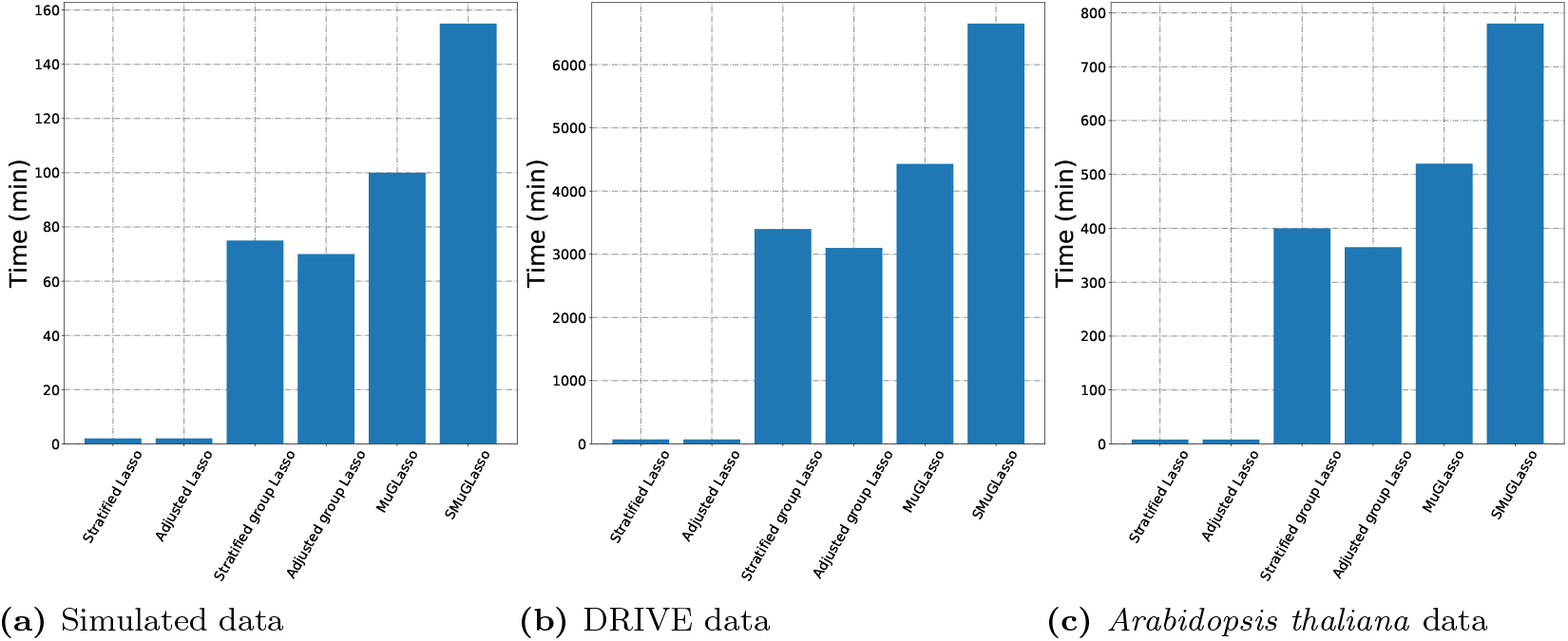
Runtimes of Lasso approaches for simulated, DRIVE and *Arabidopsis thaliana* datasets

### MuGLasso and SMuGLasso point to genes of interest not identified by classical GWAS

#### Genes identified by physical mapping annotation of SNPs selected in the DRIVE data

The breast cancer risk genes identified by physical mapping of the SNPs selected by adjusted GWAS, SMuGLasso and MuGLasso on DRIVE are detailed in Table S8 Table.

SMuGLasso and MuGLasso both recover the 9 risk genes identified by classical GWAS. SMuGLasso identifies 27 more risk genes. Of those, 17 have been previously identified in a meta-GWAS analysis containing the DRIVE data (see S10 Table), and another 8 have been found to be associated with breast cancer risk in other studies, leaving 2 genes with no previous evidence supporting their relation with the disease (see S11 Table).

MuGLasso selects the same genes as SMuGLasso, and an additional 5 genes. We have found in the literature evidence of the association with breast cancer of only 2 of those, leaving another 3 genes with no previous supporting evidence.

To summarize these findings, the distribution of evidence types for genes identified by each method is illustrated in Figure 3.

**Fig 3.**
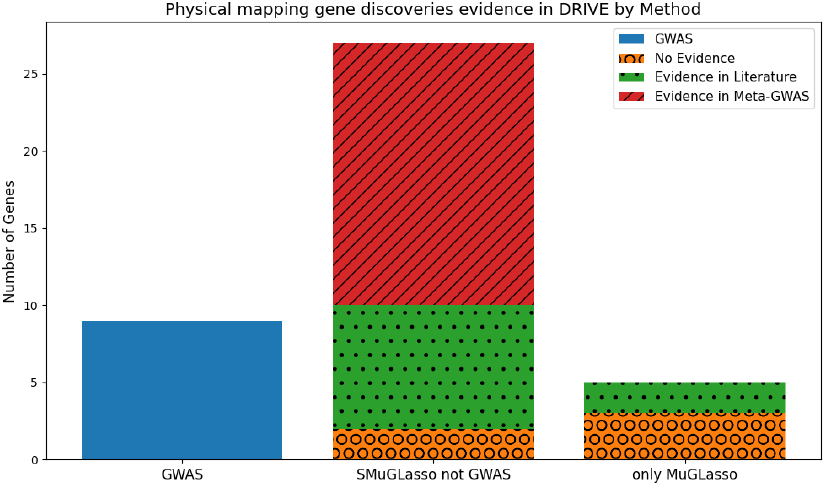
On DRIVE and through physical mapping annotation: stacked bar plot presenting the number of genes identified by different methods on the x-axis (GWAS genes, shared SMuGLasso and MuGLasso genes not identified by GWAS and only MuGLasso genes not identified by SMuGLAsso and GWAS), and categorized by evidence type (No evidence, evidence in literature and evidence in Meta-GWAS). Note that MuGLasso finds all genes selected by SMuGLasso, and SMuGLasso finds all genes selected by a classical GWAS.

#### Genes identified by physical mapping annotation of SNPs selected in the *Arabidopsis thaliana* data

The genes associated with the *Arabidopsis thaliana* DTF3 phenotype according to adjusted GWAS, SMuGLasso and MuGLasso are presented in S13 Table. Again, SMuGLasso and MuGLasso both recover all the risk genes (7 in total) identified by classical GWAS. SMuGLasso identifies an additional 41 genes, including 8 population-specific findings, and MuGLasso identifies 7 more genes on top of those selected by SMuGLasso. Only 4 of the 55 of the genes selected by MuGLasso are population-specific.

#### Genes identified by eQTL mapping annotation of SNPs selected in the DRIVE data

In addition to physical mapping, our use of eQTL functional annotations aims to uncover supplementary information about the genetic basis of breast cancer disease in DRIVE. S9 Table presents the genes obtained by both physical and eQTL mapping of the loci identified by the adjusted GWAS, SMuGLasso and MuGLasso. Using eQTL mapping adds 25 genes to the 9 identified by physical mapping of the adjusted GWAS results. 2 of those (PTHLH and TNRC6B) had been identified through the physical mapping of loci selected by SMuGLasso (and hence MuGLasso) but not the classical GWAS.

Using eQTL mapping also adds 30 genes to the list of those selected by SMuGLasso but not the classical GWAS. Of these, 26 are confirmed by the literature, as presented in S12 Table). Finally, the loci selected only by MuGLasso point to an additional 12 genes through eQTL mapping, only 2 of which are linked to breast cancer in the literature. To provide further clarification, the distribution of evidence types for these gene discoveries is illustrated on Figure 4.

**Fig 4.**
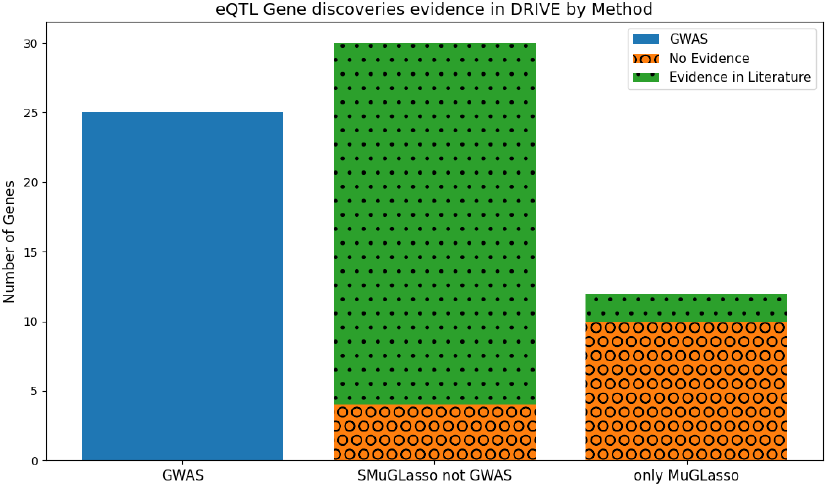
On DRIVE and through eQTL mapping annotation: stacked bar plot presenting the number of genes identified by different methods in x-axis (GWAS genes, shared SMuGLasso and MuGLasso genes not identified by GWAS and only MuGLasso genes not identified by SMuGLasso and GWAS) and categorized by evidence type (No evidence or evidence in the literature).

**Fig 5.**
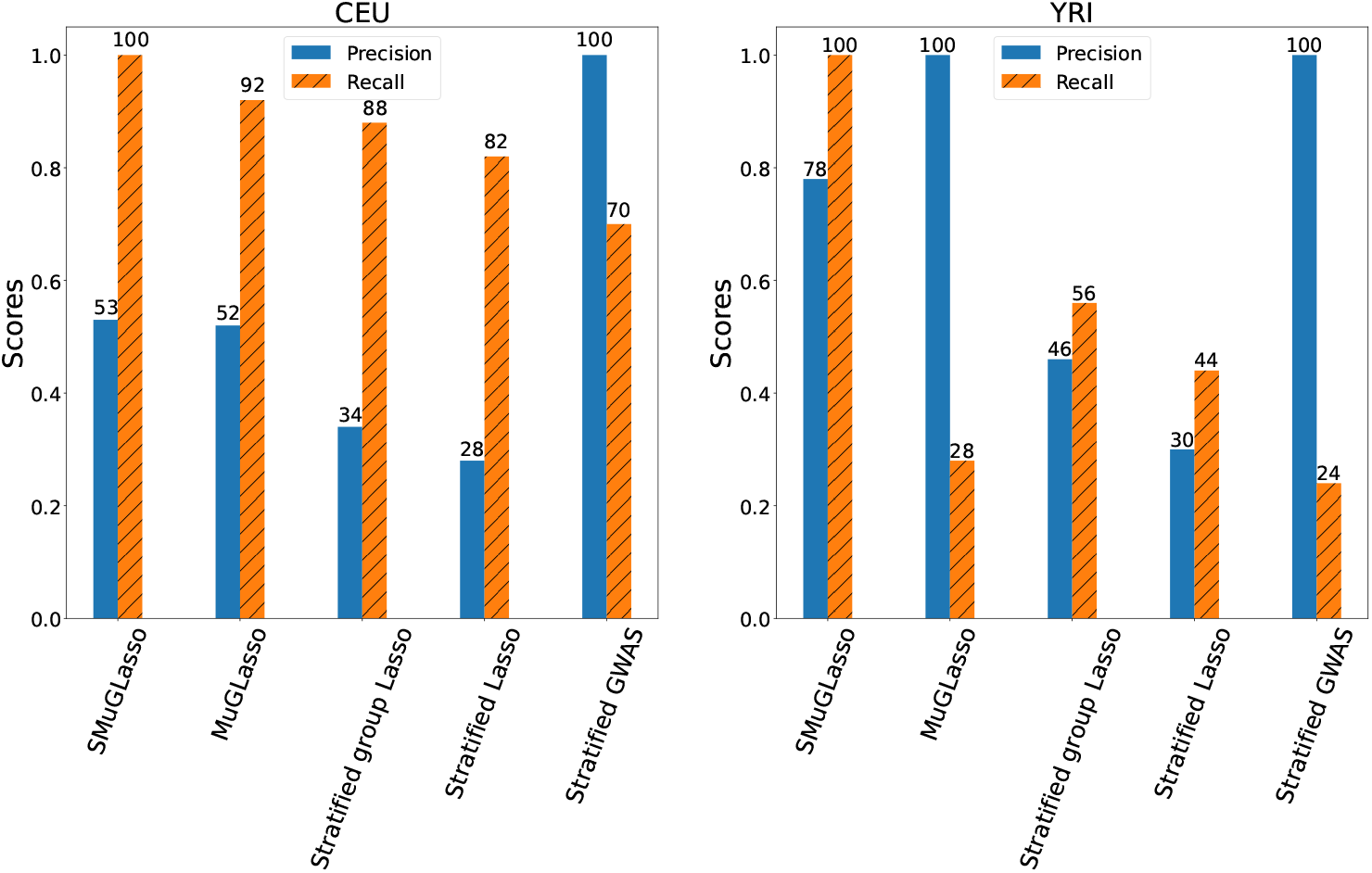
For simulated data, precision and recall of SMuGLasso, MuGLasso and the stratified approaches on the populations-specific SNPs

### SMuGLasso and MuGLasso outperform the other methods in terms of stability

Tables 4, Table 5 and S7 Table show the performance of the tested methods concerning the stability, measured by the stability index alongside the number of selected LD-groups and SNPs, along with their selection level (LD-groups or Single-SNP) respectively for simulated, DRIVE and *Arabidopsis thaliana* datasets. We use 100 subsamples to perform stability selection [12]. Indeed, the obtained metrics highlight that stability selection increases the robustness of SMuGLasso, MuGLasso and Adjusted group Lasso for the three datasets.

**Table 4.**
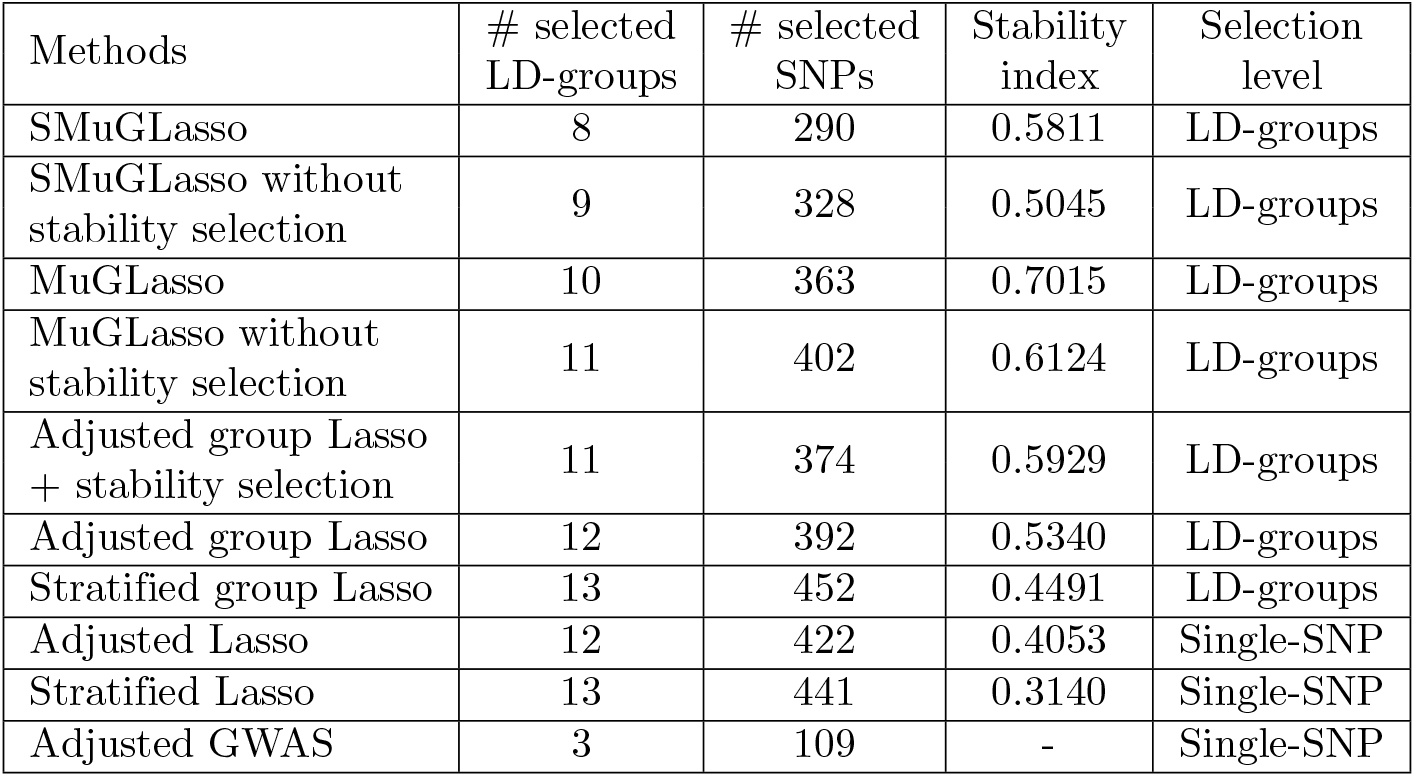
Stability index and number of selected features for different methods, on simulated data.

| Methods | # selected LD-groups | # selected SNPs | Stability index | Selection level |
| --- | --- | --- | --- | --- |
| SMuGLasso | 8 | 290 | 0.5811 | LD-groups |
| SMuGLasso without stability selection | 9 | 328 | 0.5045 | LD-groups |
| MuGLasso | 10 | 363 | 0.7015 | LD-groups |
| MuGLasso without stability selection | 11 | 402 | 0.6124 | LD-groups |
| Adjusted group Lasso + stability selection | 11 | 374 | 0.5929 | LD-groups |
| Adjusted group Lasso | 12 | 392 | 0.5340 | LD-groups |
| Stratified group Lasso | 13 | 452 | 0.4491 | LD-groups |
| Adjusted Lasso | 12 | 422 | 0.4053 | Single-SNP |
| Stratified Lasso | 13 | 441 | 0.3140 | Single-SNP |
| Adjusted GWAS | 3 | 109 | - | Single-SNP |

**Table 5.**
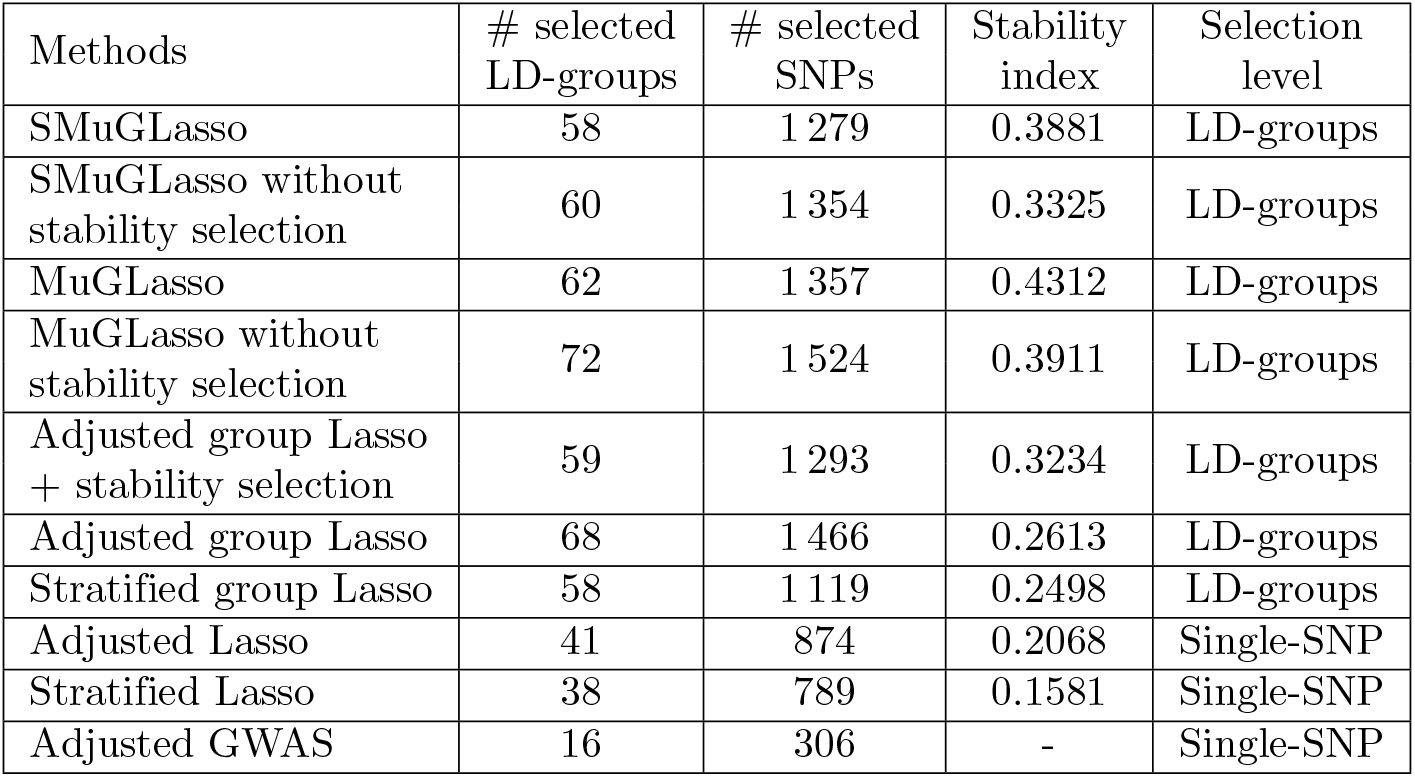
Stability index and number of selected features for different methods, on the DRIVE data set.

| Methods | # selected LD-groups | # selected SNPs | Stability index | Selection level |
| --- | --- | --- | --- | --- |
| SMuGLasso | 58 | 1 279 | 0.3881 | LD-groups |
| SMuGLasso without stability selection | 60 | 1 354 | 0.3325 | LD-groups |
| MuGLasso | 62 | 1 357 | 0.4312 | LD-groups |
| MuGLasso without stability selection | 72 | 1 524 | 0.3911 | LD-groups |
| Adjusted group Lasso + stability selection | 59 | 1 293 | 0.3234 | LD-groups |
| Adjusted group Lasso | 68 | 1 466 | 0.2613 | LD-groups |
| Stratified group Lasso | 58 | 1 119 | 0.2498 | LD-groups |
| Adjusted Lasso | 41 | 874 | 0.2068 | Single-SNP |
| Stratified Lasso | 38 | 789 | 0.1581 | Single-SNP |
| Adjusted GWAS | 16 | 306 | - | Single-SNP |

**Table 6.** Number of selected LD-groups/SNPs, across and per population, for the three data sets, for both SMuGLasso and MuGLasso.

| Data | Population | # selected LD-groups (and SNPs) |  |
| --- | --- | --- | --- |
|  |  | SMuGLasso | MuGLasso |
| Simulated data | CEU | 2 (104 SNPs) | 2 (88 SNPs) |
|  | YRI | 3 (64 SNPs) | 1 (14 SNPs) |
|  | shared (CEU and YRI) | 3 (122 SNPs) | 6 (261 SNPs) |
| DRIVE | POP1 | 5 (155 SNPs) | 6 (148 SNPs) |
|  | POP2 | 1 (21 SNPs) | 2 (43 SNPs) |
|  | shared (POP1 and POP2) | 52 (1 103 SNPs) | 54 (1 166 SNPs) |
| *A. thaliana* | POP1 | 3 (247 SNPs) | 2 (164 SNPs) |
|  | POP2 | 5 (381 SNPs) | 4 (303 SNPs) |
|  | POP3 | 1 (81 SNPs) | 0 |
|  | POP4 | 3 (232 SNPs) | 3 (232 SNPs) |
|  | POP5 | 1 (72 SNPs) | 0 |
|  | shared (5 populations) | 67 (5 354 SNPs) | 95 (7 555 SNPs) |

For the simulated data (Table 4), SMuGLasso and MuGLasso exhibit noteworthy stability, surpassing other methods. SMuGLasso selects 8 LD-groups and 290 SNPs, demonstrating a stability index of 0.5811. Similarly, MuGLasso selects 10 LD-groups and 363 SNPs with a higher stability index of 0.7015. Even without stability selection, both methods marginally increase the number of selected LD-groups and SNPs compared to classical approaches while maintaining relatively high stability indices.

For the DRIVE dataset (Table 5), SMuGLasso and MuGLasso continue to exhibit better stability than their comparison partners. SMuGLasso selects 58 LD-groups and 1,279 SNPs with a stability index of 0.3881, while MuGLasso selects 62 LD-groups and 1,357 SNPs with a stability index of 0.4312.

For the *Arabidopsis thaliana* dataset (see ), SMuGLasso and MuGLasso present once again the best stability indices. SMuGLasso selects 80 LD-groups and 6,367 SNPs with a stability index of 0.4315, while MuGLasso selects 104 LD-groups and 8,254 SNPs with a stability index of 0.5733. Thus, SMuGLasso and MuGLasso demonstrate good performance even with datasets containing relatively few samples.

Note that for methods offering selection at the single-SNP level, once a SNP is selected, we consider that the entire LD-group is selected.

MuGLasso remains the model that gives the best stability values on all datasets, followed by SMuGLasso, which outperforms the other applied feature selection methods. It’s noteworthy that SMuGLasso produces fewer selected SNPs and LD-groups compared to MuGLasso. Indeed, enforcing an additional penalty yields a sparser model, at the expense of stability; this behavior is on par with what is usually observed with lasso vs elastic net regularization.

### SMuGLasso and MuGLasso select both population-specific and shared LD-groups

SMuGLasso ensures the selection of both shared (across tasks) and task-specific LD-groups. MuGLasso can also provide such a selection at the cost of a post-processing step, which consists of removing the groups with near-zero regression coefficients for a specific task.

Table 6 presents the number of shared and population-specific LD-groups (along with the corresponding number of SNPs) selected respectively by SMuGLasso and MuGLasso on the simulated, DRIVE, and *Arabidopsis thaliana* data sets. This underscores the ability of both methods to capture task-specific genetic features while also identifying shared patterns across different populations.

For comparison, feature selection in stratified models is conducted separately for each task. Thus, the populations-specific LD-groups in stratified models correspond to LD-groups that were only selected in one population. Notably, the adjusted methods for population stratification (Adjusted group Lasso, Adjusted Lasso, and Adjusted GWAS) do not allow the selection of population-specific LD-groups.

These findings have implications for the practical application of the methods. For instance, SMuGLasso has better recall for population-specific SNPs, as illustrated by Figure 5. This figure shows the precision and recall of SMuGLasso, MuGLasso, and the stratified approaches on the population-specific SNPs, highlighting the improved performance of SMuGLasso in reducing the number of falsely selected SNPs, thanks to its additional 𝓁_1_-norm regularization.

### Candidate genes in DRIVE show breast cancer related expression

#### Gene set enrichment analysis

To further explore the usefulness of SMuGLasso, we investigated the genes it selected on the DRIVE dataset by performing gene set enrichment analyses.

S14 Table shows the top 10 pathways and processes enriched in genes selected by SMuGLasso on DRIVE. These gene sets reveal ontology terms related to biological processes and pathways implicated in breast cancer development and progression, as supported by the literature. For instance, “mammary gland morphogenosis” and “intracellular signaling by second messengers” highlight key pathways involved in breast development and cellular signaling [35, 36]. Furthermore, terms such as “ectoderm differentiation” and “endoderm differentiation” point to the importance of cellular differentiation processes in breast tissue homeostasis and tumor formation [37]. Hence, the enrichment of SMuGLasso genes in these pathways underscores their potential roles as regulators in breast cancer biology.

S15 Table, S16 Table, S17 Table, S18 Table present further enrichment analyses against various ontologies. Many of the DisGeNET disease terms that are significantly (corrected p-values *<* 0.05) enriched in genes selected by SMuGLasso (see S15 Table) correspond to subtypes of breast cancer (estrogen receptor-positive breast cancer, luminal A breast carcinoma, luminal B breast carcinoma, estrogen receptor-negative breast cancer, stage 0 breast carcinoma, mammary neoplasms). Several other enriched disease terms pertain to related diseases (uterine fibroids, squamous cell carcinoma of lung). Finally, the “breast size” trait could indeed be associated with breast cancer risk through breast tissue density, which contributes to an increased risk of developing breast cancer [38].

One cell type signature is significantly (corrected p-value *<* 0.05) enriched in genes selected by SMuGLasso: fetal thymic epithelial cells (see S16 Table). While the relationship with breast cancer genes is not immediately obvious, this could be related to the role of thymic function in mammary gland development and tumorigenesis [39].

Finally, although not significant after correction for multiple hypothesis testing, transcription factor target enrichment analysis shows enrichment of targets of the Brn-2 transcription factor (see S18 Table). These targets (FGFR2, PTLH, ELL, and ZMIZ1) have already been identified through meta-GWAS of breast cancer (see S10 Table). In addition, this is consistent with findings demonstrating that this transcription factor promotes invasion and metastasis in triple-negative breast cancer cells [40].

#### Protein-protein interaction analysis

We also used Metascape to identify the modules of the protein-protein interaction network formed from known interactions between genes identified through physical mapping of the SNPs selected by the Adjusted GWAS approach, SMuGLasso and MuGLasso, which are shown on Figure 6. The modules obtained when adding genes identified through eQTL mapping can be visualized on Figure . Pathway and process enrichment analysis of the genes in the two modules identified by SMuGLasso (Figure 6c) highlights three significant processes, described in Table 7. These highlights the ability of SMuGLasso to identify more relevant disease genes than a classical GWAS approach. Indeed, as breast cancer often originates from aberrant growth and dysfunction within mammary gland structures, mammary gland morphogenesis processes may serve as crucial drivers or modifiers of tumor initiation and progression [35, 41]. Furthermore, phosphorylation is well-known to play a role in regulating cellular processes such as proliferation, migration, and survival, all of which are dysregulated in cancer [42, 43] in general and in breast cancer in particular [44].

**Table 7.** Pathway and process enrichment analysis of the modules of the PPI of known interactions between genes identified through physical mapping of the SNPs selected by SMuGLasso on DRIVE.

| GO | Description | Log10(P) | Gene Hits |
| --- | --- | --- | --- |
| GO:0060443 | mammary gland morphogenesis | -6.4 | ESR1, FGFR2, TGFBR2 |
| GO:0016310 | phosphorylation | -5.5 | FGFR2, HK1, MAP3K1, TGFBR2, NEK10 |
| GO:0022612 | gland morphogenesis | -5.2 | ESR1, FGFR2, TGFBR2 |

**Fig 6.**
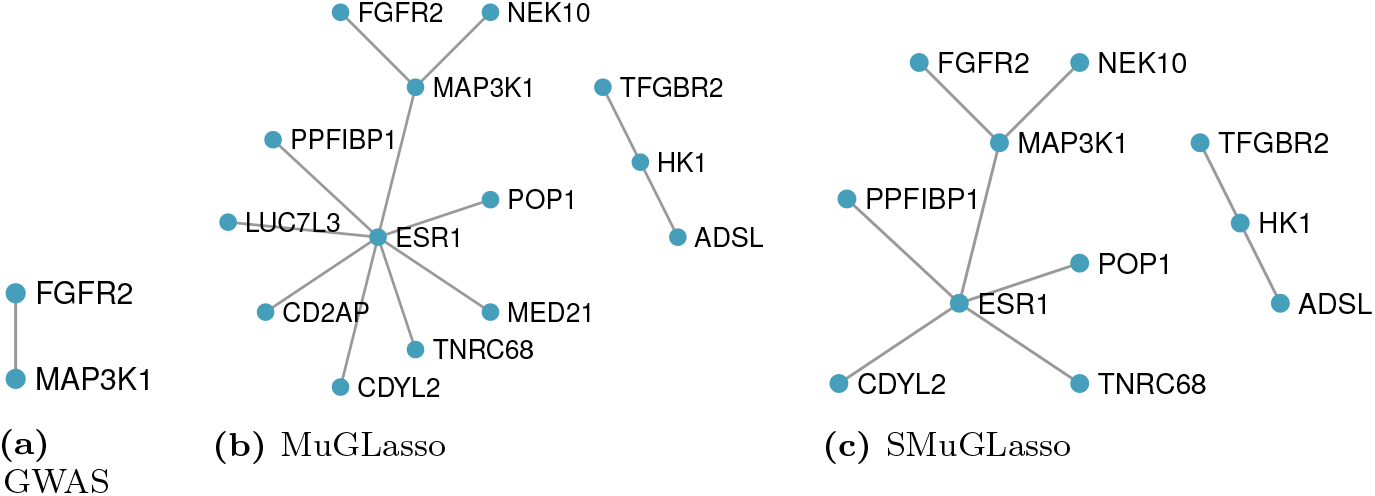
Modules of the PPI of known interactions between genes identified through physical mapping of the SNPs selected by Adjusted GWAS, SMuGLasso and MuGLasso on DRIVE.

All in all, gene set enrichment analysis of the genes pinpointed by SMuGLasso highlight the relevance of the disease genes it detects in addition to a classical GWAS, suggesting that this tool can be used to provide valuable insights into the molecular mechanisms underlying phenotypes.

#### Quantitative pathway enrichment analysis comparison

To compare pathway enrichments between Adjusted GWAS, MuGLasso and SMuGLasso, we conduct a quantitative analysis to assess their biological significance. We extract the Z-scores for each common pathway across the three methods (Adjusted GWAS, MuGLasso and SMuGLasso). For this analysis, we consider the top 50 pathways with the highest Z-scores for each method and then focus on the common pathways among these top-ranked sets. For each shared pathway, we compute the Z-score ratio for each pairwise comparison of methods (MuGLasso/GWAS to assess MuGLasso vs. Adjusted GWAS, MuGLasso/Adjusted GWAS to assess Adjusted GWAS vs. SMuGLasso and SMuGLasso/MuGLasso to assess MuGLasso vs. SMuGLasso). Thus, a Z-score ratio higher than 1 indicates that the first method in the pair (e.g., MuGLasso in the MuGLasso/Adjusted GWAS comparison) shows greater enrichment for that pathway than the second method (e.g., Adjusted GWAS). Conversely, a Z-score ratio of less than 1 indicates that the second method in the pair shows greater enrichment for that pathway compared to the first method. We further assess the significance of differences in pathway enrichments using paired t-tests.

In Figure 7, the box plots illustrate comparisons of pathway enrichments between SMuGLasso, Adjusted GWAS, and MuGLasso. Notably, SMuGLasso shows higher pathway and process enrichment compared to both Adjusted GWAS and MuGLasso, as indicated by the higher Z-score ratios. Furthermore, the performed paired t-tests reveal statistically significant differences between SMuGLasso and MuGLasso, confirming that SMuGLasso identifies pathways with greater biological significance compared to MuGLasso.

**Fig 7.**
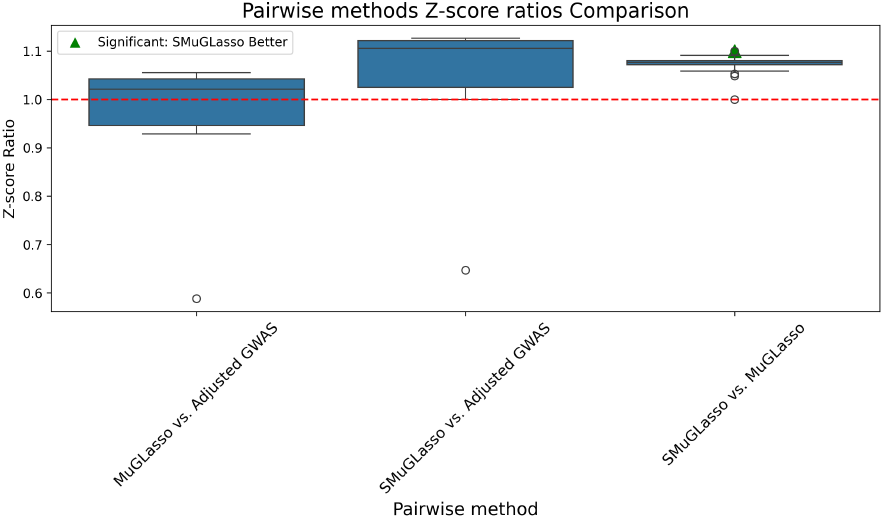
On DRIVE, box plots representing the distribution of Z-score ratios for gene sets enrichments across three pairwise comparisons: MuGLasso vs. Adjusted GWAS, SMuGLasso vs. Adjusted GWAS, and SMuGLasso vs. MuGLasso. The Z-score ratio is computed for each shared pathway between the methods. The green triangle indicates significant differences in enrichment (p-value *<* 0.005) as determined by paired t-tests. The red dashed line at *y* = 1 represents equal enrichment by both methods.

Further figures illustrate that SMuGLasso exhibits greater enrichment than MuGLasso across all gene sets (S22 Fig), including DisGeNET gene sets (S19 Fig). We emphasize the DisGeNET gene sets in these analysis, as they are the only ones for which classical GWAS shows enrichment. Compared to GWAS, SMuGLasso demonstrates higher enrichment in 4 out of 6 pathways, equivalent enrichment in 1 pathway, and lower enrichment in 1 other pathway (see S20 Fig), whereas MuGLasso shows better enrichment than GWAS in 3 pathways, equal enrichment in 1 pathway, and lower enrichment in 2 pathways (S21 Fig).

Notably, the top enrichment results determined through pathway analysis are similar regardless of whether or not eQTL gene lists are included.

### Candidate genes in *Arabidopsis thaliana* show flowering time related expression

We present in S13 Table the list of mapped genes using TAIR10 gff3 mapping of SNPs selected on the *Arabidopsis thaliana* data set by Adjusted GWAS, SMuGLasso and MuGLasso. We conducted pathway enrichment analysis and observed distinct differences in enrichments among the tested methods. SMuGLasso identified 48 genes, among which two pathways are significantly overrepresented: gravitropism and response to carbohydrate. MuGLasso discovered 55 genes, among which only the gravitropism pathway is overrepresented. By contrast, Adjusted GWAS identified 7 genes, for which the enrichment analysis did not yield any pathway.

An advantage of SMuGLasso compared to MuGLasso is that it finds more pathways with fewer gene discoveries, suggesting it is likely more efficient. The pathways identified by SMuGLasso and MuGLasso, namely gravitropism and response to carbohydrate, have potential relevance to the time until the first open flower phenotype. Gravitropism, the orientation or growth of plants in response to gravity, could influence floral development by affecting how plants orient their growth and allocate resources [45]. The response to carbohydrate pathway discovered by SMuGLasso may also be significant, as carbohydrates are crucial for energy storage and signaling, which can impact plant growth and development, including flowering time [46]. These observations suggest that SMuGLasso and MuGLasso methods are more effective in uncovering biologically relevant pathways that could explain variations in the flowering time phenotype compared to Adjusted GWAS, which did not identify any related pathways.

Finally, Figure 8 shows the modules of the protein-protein interaction network of known interaction between the genes identified by SMuGLasso and MuGLasso. The MuGLasso PPI network appears denser and more informative, with additional interactions and nodes such as FUS9 and ICK1, compared to the SMuGLasso PPI network, which is sparser and has fewer connections. However, upon further investigation, we did not find any evidence linking these nodes, particularly FUS9 and ICK1, to flowering time regulation in *Arabidopsis thaliana*.

**Fig 8.**
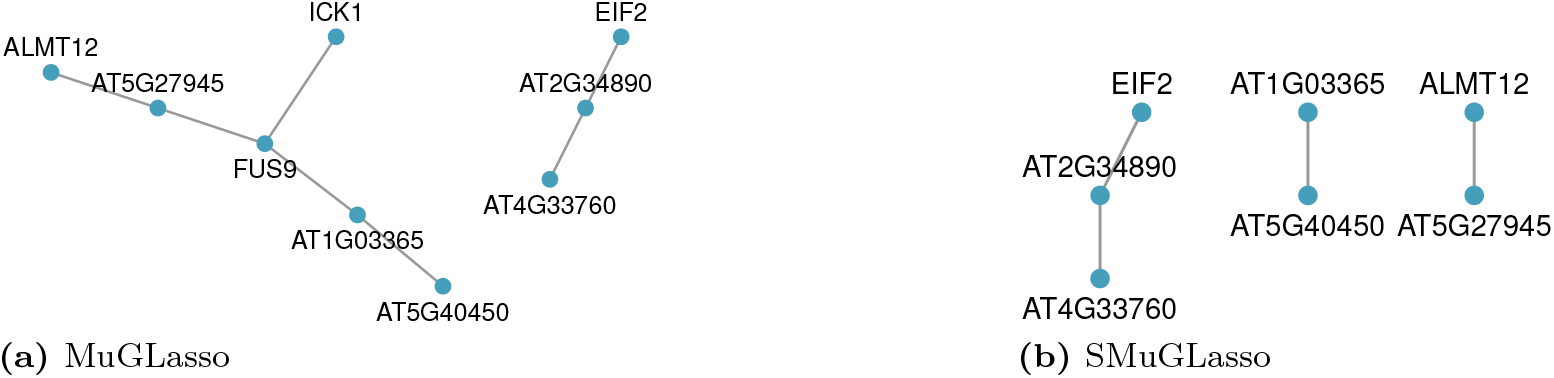
Modules of the PPI of known interactions between genes identified through physical mapping of the SNPs selected by MuGLasso and SMuGLasso on the *Arabidopsis thaliana* data set.

## Discussion

We have presented in this paper SMuGLasso, an extension of MuGLasso for the identification of relevant SNPs from GWAS data across multiple populations. The proposed model is based on a multitask framework in which the tasks are genetic populations and features are clustered in groups. The selection is performed at the scale of LD-groups. The populations are identified using PCA and k-means to assign each sample to a subpopulation. This setting alleviates the curse of dimensionality and addresses population stratification in diverse populations. Compared to MuGLasso, SMuGLasso includes an additional regularization term which enforces task-specific sparsity at the level of LD-groups. Thus, our model provides indeed a more precise recovery of risk regions related to the phenotype at the population-specific level.

Our simulations demonstrate that SMuGLasso outperforms MuGLasso and other methods in accurately identifying population-specific disease loci, while also minimizing potential false discoveries. While MuGLasso shows commendable stability, SMuGLasso closely follows, exhibiting robust stability indexes across various datasets with a reduced number of selected LD-groups/SNPs. The application of stability selection techniques further bolsters SMuGLasso’s reliability in terms of stability measurements.

A significant advancement in our study is addressing the computational challenges posed by the additional penalty in MuGLasso, achieved through the implementation of gap-safe screening rules. This ensures efficient processing for both qualitative and quantitative phenotypes.

Lastly, we have detailed the genes identified by both SMuGLasso and MuGLasso in our real data analyses, and we performed pathway analysis with biological interpretation for the entire gene lists. Interestingly, SMuGLasso’s findings are more consistent with the literature than those that are specific to MuGLasso. However, we encountered limitations in investigating pathway analysis specific to populations due to the absence of tools that adequately consider population structure. Despite our efforts, we were unable to find evidence in our findings for pathway enrichment specific to particular populations. Looking ahead, our goal is to delve into pathway analysis to unravel the biological mechanisms underpinning the phenotypes of interest in diverse population studies, as revealed by the identified risk genes.

Despite the implementation of gap-safe screening rules, the computational load is still significant, especially when dealing with extremely large datasets. This could limit the applicability in broader GWAS data where computational resources are a constraint. The efficacy of SMuGLasso heavily relies on the ability of PCA and k-means clustering in identifying subpopulations. Misclassification or suboptimal clustering can potentially impact the final results. One solution is to focus on enhancing the clustering stability through a hierarchical structure. Moreover, given the additional regularization term, there is a risk that the model might become biased towards tasks with more samples, potentially overlooking key insights in the less-represented tasks/populations.

Introducing a weighting scheme that balances the influence of each task, particularly giving more weight to those with fewer samples, might help in addressing the imbalance, or integrating additional external datasets to bolster the sample sizes of underrepresented tasks could be beneficial. In addition, investigating different regularization terms for the population-specific LD-groups selection or hybrid approaches could potentially improve the model’s performance in identifying disease-relevant loci. Furthermore, there remains an essential avenue for future work in rigorously integrating alternative stability selection methods, which could further improve SMuGLasso’s robustness. In conclusion, while SMuGLasso presents a novel framework in the field of GWAS analysis, especially in the precise identification of population-specific risk loci, our ongoing efforts to refine its computational efficiency, enhance clustering accuracy, and balance task representation will be essential in realizing its full potential in unraveling the complex genetic mechanisms of diseases.

## Supporting information

## S1 Appendix SMuGLasso method details

## Population structure

Diverse and admixed populations studies offer a unique, yet complex, opportunity. On the one hand, they provide an excellent means to increase the number of samples, unlike homogeneous studies where the number of samples is in most data very restricted. Indeed, genotyping hundreds of thousands of participants from different ancestries to study a phenotype of interest helps to alleviate the curse of dimensionality. On the other hand, such analyses require close attention to the confounder raised by population stratification, that is, when association is detected on the population structure rather than on the phenotype of interest. The presence of population stratification is one of the major problems in association studies as it increases type I error and leads to ambiguous results. This is particularly true when allele frequency differences in cases and controls are due to differences in ancestry rather than association between SNPs and disease. To counteract the issue, several correction methods have been developed, including genomic control, Principal Component Analysis (PCA)-based methods [47–54], and Linear mixed models [29, 55, 56]. Each of these approaches offers specific benefits and is designed to mitigate the impact of population structure on study results. However, a critical aspect to consider in these adjustment methods is the potential for overcorrection, particularly concerning causal SNPs in certain populations. Additionally, these techniques may not fully account for the presence of population-specific variants that are associated with the disease under study.

Therefore, we observe an existent need in GWAS field to develop efficient frameworks that profit from the advantages provided by diverse studies. Such frameworks should address the issues posed by population stratification, as well as consider the existence of population-specific causal LD-groups.

Consequently, to handle population structure in our model, we assign genetic subpopulations to specific input tasks. This is achieved by using PCA in conjunction with k-means clustering, which allows us to accurately determine the subpopulation to which each sample belongs.

## Linkage disequilibrium groups clustering

LD causes genetic variants to be correlated, indicating that nearby alleles are inherited together more often than expected by chance, thereby influencing the identification of disease-associated SNPs. Hence, current approaches in association studies use hierarchical clustering to construct LD-groups, effectively grouping variants based on their correlation patterns. Applying feature selection on LD-groups rather than on single SNPs improves remarkably the stability of the selection. By performing the selection at the LD-groups level, the method reduces the selection options for regularization models, and hence addresses the curse of dimensionality, improving the analysis by focusing on correlated groups of genetic variants. In practice, we have used adjacency-constrained hierarchical clustering algorithm to form the LD-groups assigned to SMuGLasso with adjclust R package [10].

## Stability selection

We use the stability selection procedure developed by [12] where they propose improving the stability using a subsampling method. In this formulation, variable selection is performed repeatedly on subsamples. The subsampling approach can be used to determine the amount of regularization needed to control the familywise error type I rate. Stability selection is a feature selection based method, it can be combined with several existing methods and aims to improve their performance. The procedure relies on computing the stability path, which represents the probability of a feature to be selected across random subsamples, as a function of the regularization parameter.

We denote by *I* a random subsample of {1, …, *n*} of size ⌊*n/*2⌋, we call 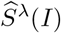 the set of features selected by the selection procedure of interest (for example, Lasso), with a hyperparameter *λ*, on this subsample of the data. For any feature *j* ∈ {1, …, *p*}, we call 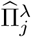 the probability that feature *j* is selected on a random subsample of size ⌊*n/*2⌋ of the data. This probability is determined, given *m* such random subsamples *I*_1_, *I*_2_, …, *I*_*m*_, as the proportion of those subsamples for which the feature selection procedure selects a feature *j*:

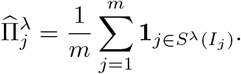

Finally, given a threshold 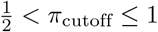 (in this work, we used *π*_cutoff_ = 0.75), the stable set of selected features is determined as:

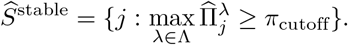

We detail below the Theorem 1 used to bound the number of expected false selected features:

## Theorem 1: Stability selection

Let’s assume the set of features with non-zero coefficients by *S* = {*j* : *β*_*j*_ ≠ 0}, and the set of features with zero coefficients by *N* = {*j* : *β*_*j*_ = 0}. Assuming that the distribution of 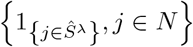 is exchangeable for all *λ* ∈ ℝ^+^. Also, assuming that the original procedure is not worse than the random setting, i.e. for any *λ* ∈ ℝ^+^:

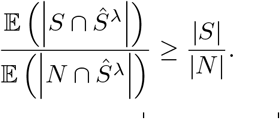

The number of falsely selected variables *V* = | *N* ∩ Ŝ^stable^ | is then bounded by:

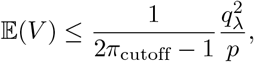

where *q*_*λ*_ = 𝔼 (|*S*_*λ*_(*I*)|) denotes the expected number of selected variables. The desired calibration is to obtain 𝔼(*V* ) ≤ *α* with *α* small.

**The stability of the selection measurement** The method is carried out with the Pearson similarity index [57] and represents an extension of [58]. Let’s assume 𝒵 = {*s*_1_, …, *s*_*M*_ } is the set of *M* selected features, where each *s*_*u*_ is a subset of the features. The total number of features is denoted by *p* and the number of features selected on the *u*^*th*^ feature set is given by *k*_*u*_. A set of selected features *s*_*u*_ can be represented by an indicator vector *z*_*u*,._ ∈ {0, 1}^*p*^, where *z*_*u*,*j*_ = 1 if feature *j* is selected and 0 otherwise. The Pearson correlation between two feature sets *s*_*u*_ and *s*_*v*_ is presented by the following equation:

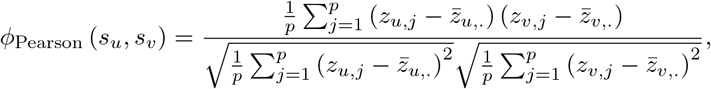

where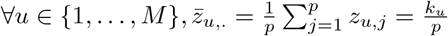

The stability of the selection based on Pearson correlation can be rewritten as follows:

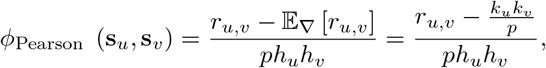

where ∀*u* ∈ {1, …, *M* }, 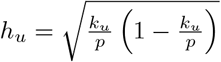 and *r*_*u*,*v*_ denotes the number of features in common between the feature sets *s*_*u*_ and *s*_*v*_. 𝔼_∇_ is an adjustment term equal to the expected value of *r*_*u*,*v*_ when the feature selection model selects randomly *k*_*u*_ and *k*_*j*_ features from all features *p*.

When the number of selected features *k* is the same for all feature sets, assuming *S* an index of the variability in the choice of features, the Pearson correlation is established as follows:

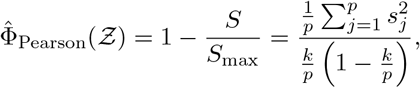

where *S*_max_ is the maximal value of *S* when the feature selection model selects *k* features per feature set. 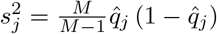 corresponds to the sample variance of selection of the *j*^*th*^ feature. 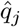 corresponds to the observed frequency of the selection of a feature *j*, as well as the sample mean of the variable Z_*j*_.

## S2 Appendix. DRIVE

## Data access

The dataset “General Research Use” in DRIVE Breast Cancer OncoArray Genotypes is available from the dbGaP controlled-access portal, under Study Accession phs001265.v1.p1 (https://www.ncbi.nlm.nih.gov/projects/gap/cgi-bin/study.cgi?study_id=phs001265.v1.p1). Researchers can gain access the data by applying to the data access committee, see https://dbgap.ncbi.nlm.nih.gov.

### Ethics approval

The dataset was obtained from NIH after ethical review of project #17707, titled “Network-guided multi-locus biomarker discovery”, and used under approval of this request (#67806-4).

## Acknowledgments

OncoArray genotyping and phenotype data harmonization for the Discovery, Biology, and Risk of Inherited Variants in Breast Cancer (DRIVE) breast-cancer case control samples was supported by X01 HG007491 and U19 CA148065 and by Cancer Research UK (C1287/A16563). Genotyping was conducted by the Center for Inherited Disease Research (CIDR), Centre for Cancer Genetic Epidemiology, University of Cambridge, and the National Cancer Institute. The following studies contributed germline DNA from breast cancer cases and controls: the Two Sister Study (2SISTER), Breast Oncology Galicia Network (BREOGAN), Copenhagen GeneralPopulation Study (CGPS), Cancer Prevention Study 2 (CPSII), The European Prospective Investigation into Cancer and Nutrition (EPIC), Melbourne Collaborative Cohort Study (MCCS), Multiethnic Cohort (MEC), NashvilleBreast Health Study (NBHS), Nurses Health Study (NHS), Nurses Health Study 2 (NHS2), Polish Breast CancerStudy (PBCS), Prostate Lung Colorectal and Ovarian Cancer Screening Trial (PLCO), Studies of Epidemiologyand Risk Factors in Cancer Heredity (SEARCH), The Sister Study (SISTER), Swedish Mammographic Cohort (SMC), Women of African Ancestry Breast Cancer Study (WAABCS), Women’s Health Initiative (WHI).

---

**S3 Fig.**
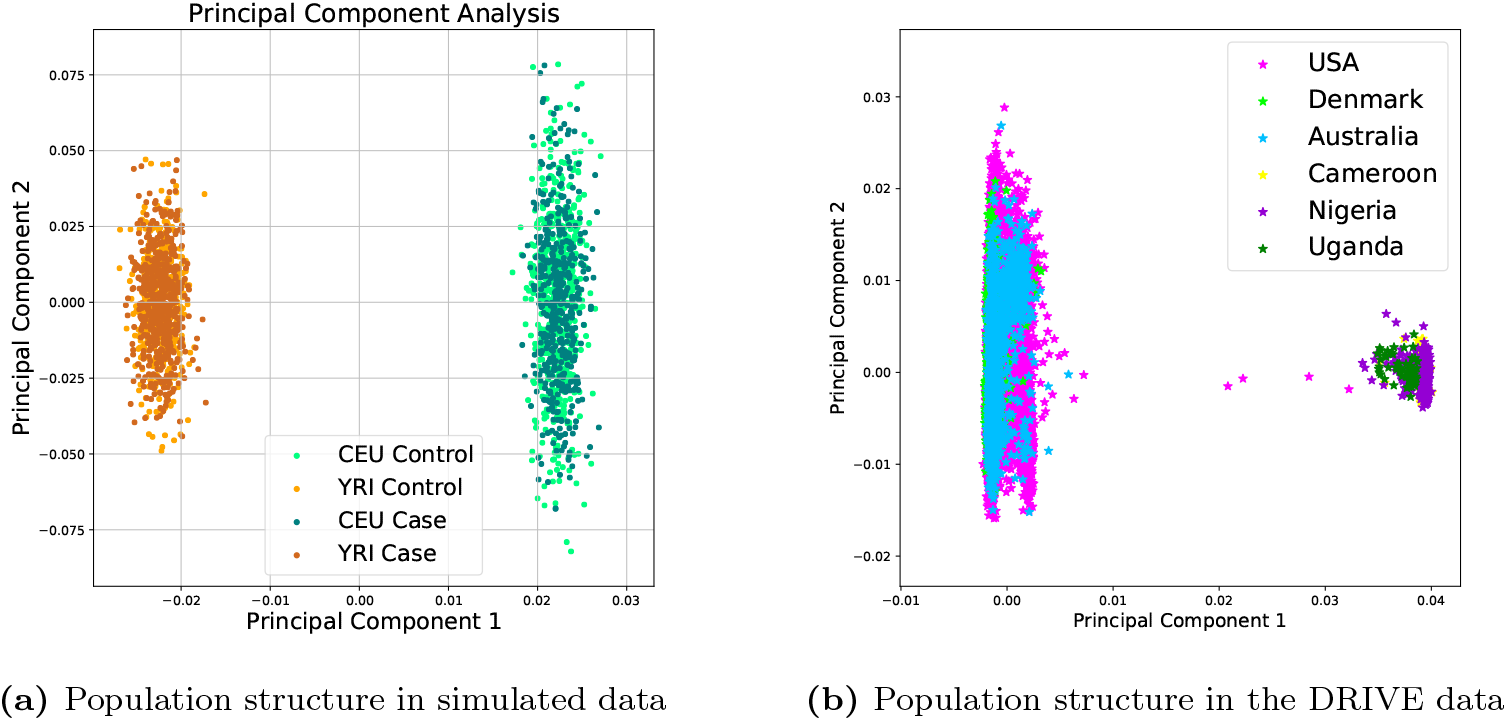
PCA for simulated and DRIVE datasets.

**S4 Fig.**
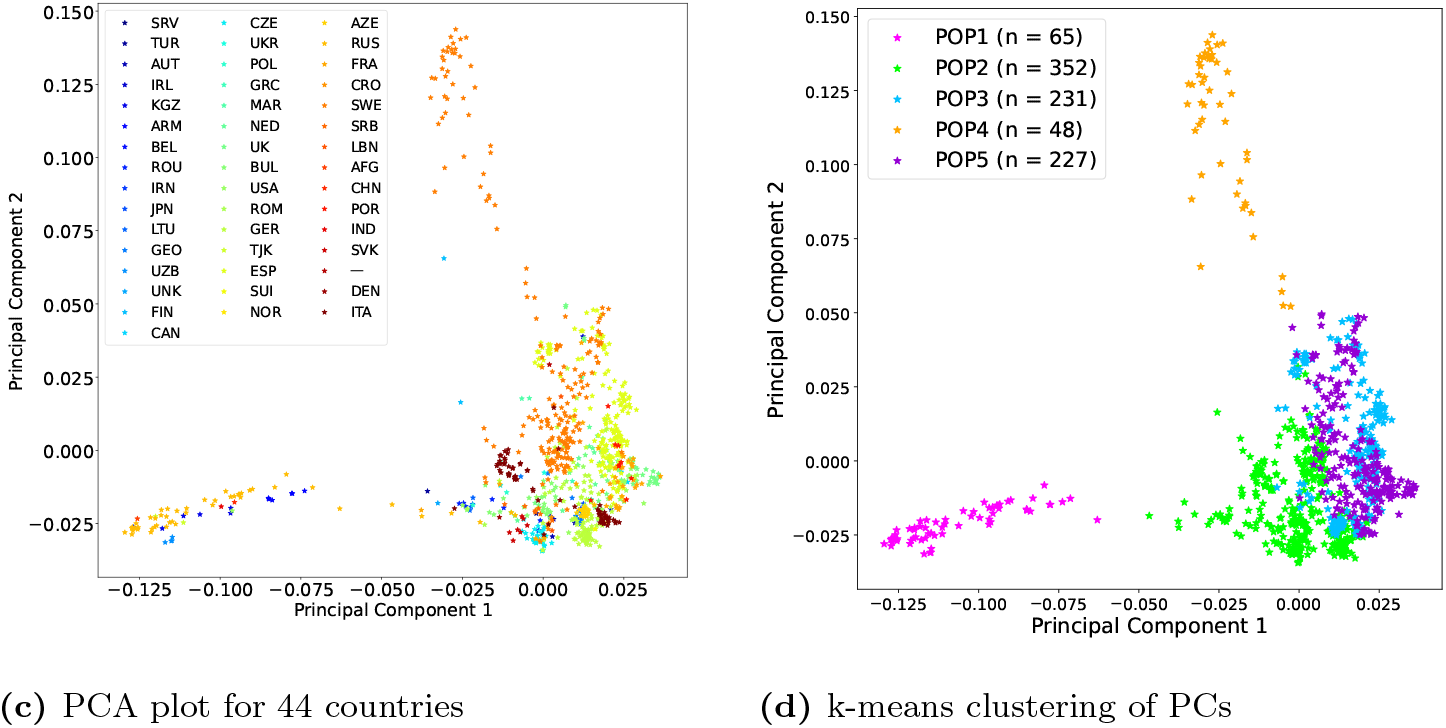
PCA for *Arabidopsis thaliana* dataset Projection of the *Arabidopsis thaliana* genotypes on the first two PCA components. Samples originate from 44 countries. We identified 5 subpopuations through K-means clustering of the data.

**S5 Fig.**
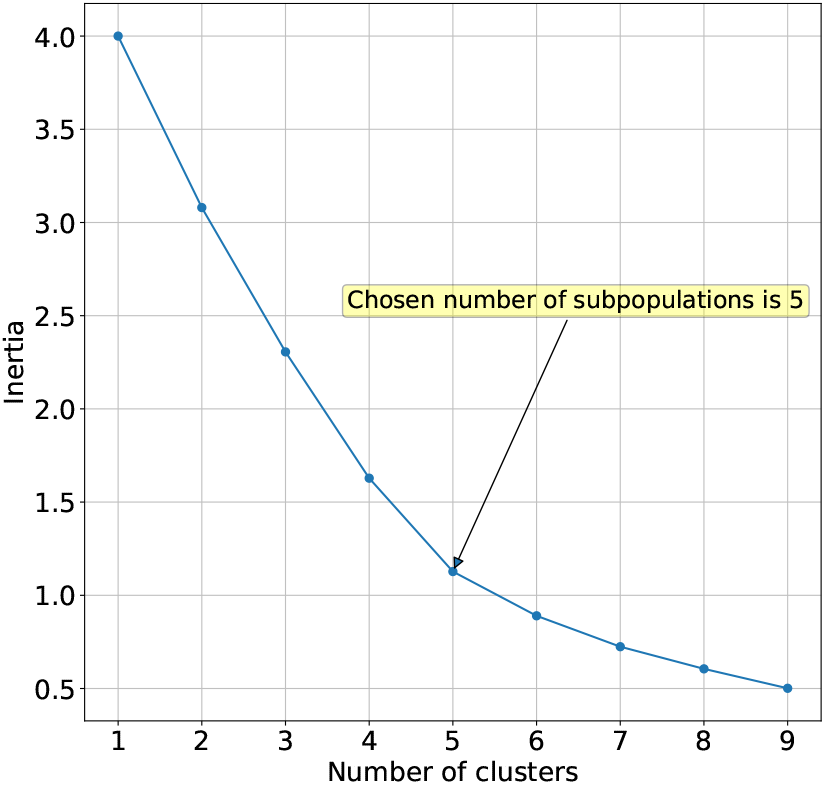
Inertia Plot for K-means Clustering of *Arabidopsis thaliana*.

**S6 Table.** Subpopulations of *Arabidopsis thaliana* with the corresponding countries and the number of samples included in each subpopulation.

| Populations | Countries (# samples) |
| --- | --- |
| POP1 | Russia(42), Armenia(7), Kyrgyzstan(5), Uzbekistan(4), Tajikistan(2), USA(1), India(1), Romania(1), China(1), Afghanistan(1) |
| POP2 | Sweden(87), Germany(84), Italy(43), Czechia(32), Bulgaria(23), Russia(8), Romania(8), UK(7), Slovakia(7), Serbia(7), Austria(6), USA(6), France(6), Lithuania(4), Switzerland(3), Netherlands(3), Poland(3), Denmark(2), Japan(2), Croatia(2), Belgium(1), Finland(1), Norway(1), Greece(1), Lebanon(1), Turkey(1), North Vietnam(1), Unknown (1) |
| POP3 | Spain(158), Italy(20), Azerbaijan(16), Georgia(14), Portugal(8), Bulgaria(5), Maroc(2), Greece(2), France(2), Ireland(1), Lebanon(1), Unknown(1) |
| POP4 | Sweden(46), Finland(1) |
| POP5 | Sweden(67), UK(43), France(32), Germany(28), USA(27), Italy(9), Spain(9), Netherlands(7), Canada(2), Ireland(1), Switzerland(1), Portugal(1) |

**S7 Table.** Stability index and number of selected features for different methods on *Arabidopsis thaliana*

| Methods | # selected LD-groups | # selected SNPs | Stability index | Selection level |
| --- | --- | --- | --- | --- |
| SMuGLasso | 80 | 6 367 | 0.4315 | LD-groups |
| SMuGLasso without stability selection | 87 | 7 220 | 0.3883 | LD-groups |
| MuGLasso | 104 | 8 254 | 0.5733 | LD-groups |
| MuGLasso without stability selection | 149 | 10 935 | 0.5040 | LD-groups |
| Adjusted group Lasso + stability selection | 90 | 6 944 | 0.4489 | LD-groups |
| Adjusted group Lasso | 114 | 8 358 | 0.3654 | LD-groups |
| Stratified group Lasso | 133 | 10 135 | 0.3147 | LD-groups |
| Adjusted Lasso | 112 | 9 258 | 0.2600 | Single-SNP |
| Stratified Lasso | 135 | 9 897 | 0.2140 | Single-SNP |
| Adjusted GWAS | 7 | 31 | - | Single-SNP |

**S8 Table.** Potential breast cancer risk genes identified through physical (within 10kb) mapping of the loci selected by Adjusted GWAS, SMuGLasso and MuGLasso. CEU-specific selected genes are highlighted in blue and YRI-specific selected genes are highlighted in red. The remaining genes (in black) are risk genes shared across all populations.

|  |  |
| --- | --- |
| Adjusted GWAS | **ITPR1, MRPS30, MAP3K1, SETD9, MIER3, EBF1, FGFR2, TOX3, MKL1.** |
| SMuGLasso | **ITPR1, MRPS30, MAP3K1, SETD9, MIER3, EBF1, FGFR2, TOX3, MKL1,** ADSL, ASTN2, CACNA1I, CCDC170, CCDC91, CDYL2, **DIRC3**, ELL, ESR1, FTO, GRHL1, HK1, **HRSP12**, KCNU1, NEK10, NUP205, PAX9, PTHLH, POP1, PPFIBP1, **REP15, SGSM3**, SSBP4, TGFBR2, TNRC6B, ZMIZ1, ZNF365. |
| MuGLasso | **ITPR1, MRPS30, MAP3K1, SETD9, MIER3, EBF1, FGFR2, TOX3, MKL1,** ADSL, ASTN2, C7orf73, CACNA1I, CCDC170, CCDC91, CCSER1, CD2AP, CDYL2, **DIRC3**, ELL, **ESR1**, FTO, GRHL1, HK1, HRSP12, KCNU1, **LUC7L3, MED21**, NEK10, NUP205, PAX9, PTHLH, POP1, PPFIBP1, **REP15, SGSM3**, SSBP4, TGFBR2, TNRC6B, ZMIZ1, ZNF365. |

**S9 Table.**
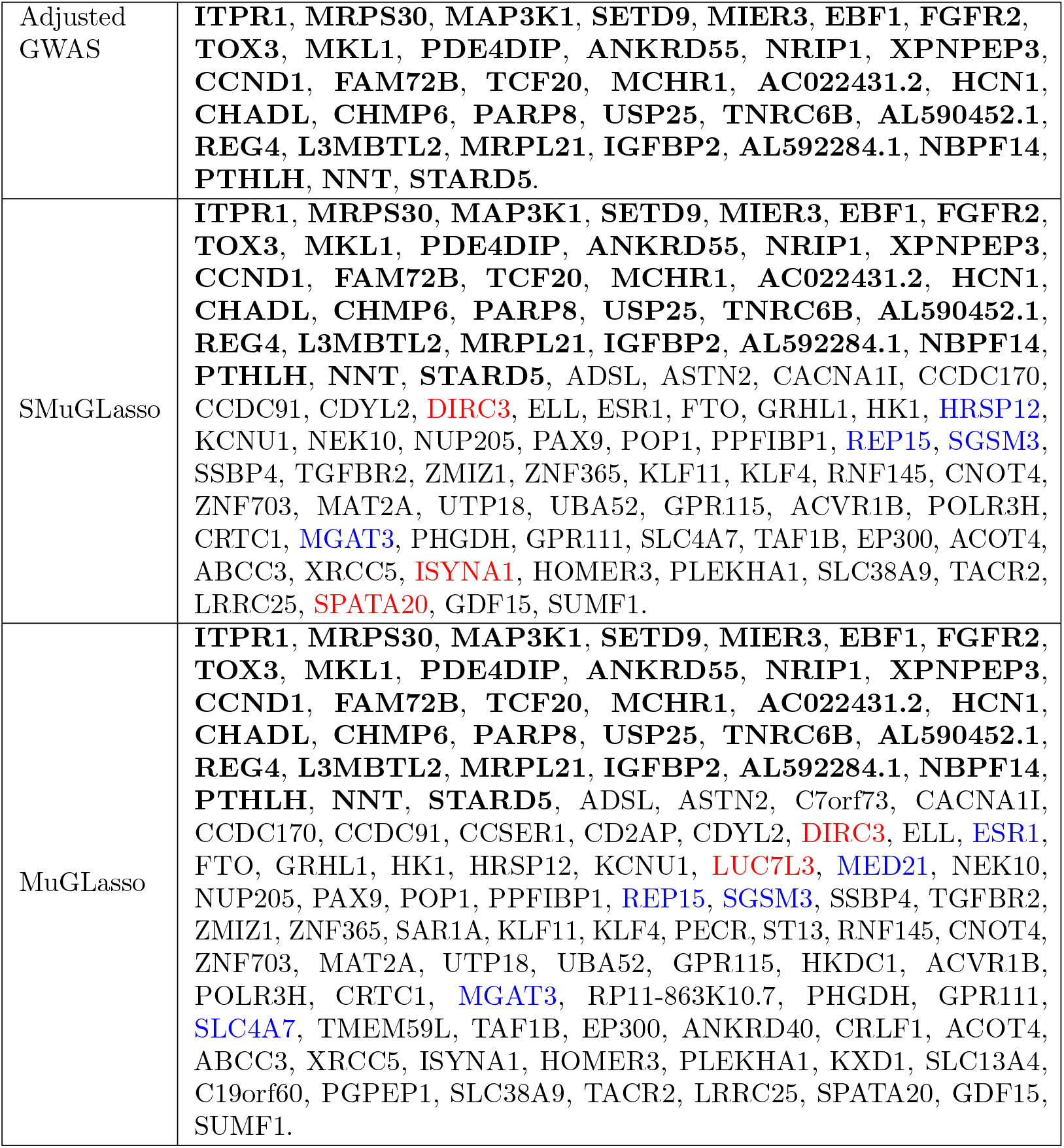
Breast cancer risk loci detected by SMuGLasso and MuGLasso on DRIVE Potential breast cancer risk genes identified through both physical (within 10kb) and eQTL mapping of the loci selected by Adjusted GWAS, SMuGLasso and MuGLasso. CEU-specific selected genes are highlighted in blue and YRI-specific selected genes are highlighted in red. The remaining genes (in black) are risk genes shared across all populations.

**S10 Table.** Potential breast cancer risk genes identified through both physical (within 10kb) and eQTL mapping of the loci selected by MuGLasso or/and SMuGLasso and not the adjusted GWAS, found in meta-GWAS including the samples used in this work.

| Gene symbols | Evidence |
| --- | --- |
| *ASTN2* | [59] |
| *CCDC170* | [59–63] |
| *CDYL2* | [59, 61, 62] |
| *DIRC3* | [59, 61–63] |
| *ELL* | [59, 61–63] |
| *ESR1* | [59, 60, 62, 63] |
| *FTO* | [59–63] |
| *GRHL1* | [59] |
| *KCNU1* | [59, 62] |
| *NEK10* | [59, 61–63] |
| *PAX9* | [59, 61, 62] |
| *PTHLH* | [59–63] |
| *SSBP4* | [59] |
| *TGFBR2* | [59, 61, 62] |
| *TNRC6B* | [59] |
| *ZMIZ1* | [59, 61, 62] |
| *ZNF365* | [59, 63] |

**S11 Table.** Potential breast cancer risk genes identified through both physical (within 10kb) and eQTL mapping of the loci selected by MuGLasso or/and SMuGLasso and not the adjusted GWAS, found to be associated with breast cancer risk or tumor growth in the literature.

| Gene symbols | Evidence |
| --- | --- |
| *ADSL* | oncogenic driver in triple negative breast cancer [64] |
| *CACNA1I* | underexpressed in breast cancer [65] |
| *CCDC91* | likely target gene of breast cancer risk variants [66] |
| *NUP205* | forms a complex with NUP93 which regulates breast tumor growth [67] |
| *POP1* | expression correlates with prognosis in breast cancer [68] |
| *PPFIBP1* | promotes cell motility and migration in breast cancer [69] |
| *SGSM3* | associated with breast cancer in a Chinese population [70] |
| *HK1* | promote breast cancer cell proliferation, migration and invasion [42] |
| *LUC7L3* | targeted by miR-370-5p, suppresses proliferation and invasion in breast cancer cells [71] |
| Other genes |  |
| *C7orf73*, *CCSER1*, *CD2AP*, *HRSP12*, *MED21*, *REP15* |  |

**S12 Table.**
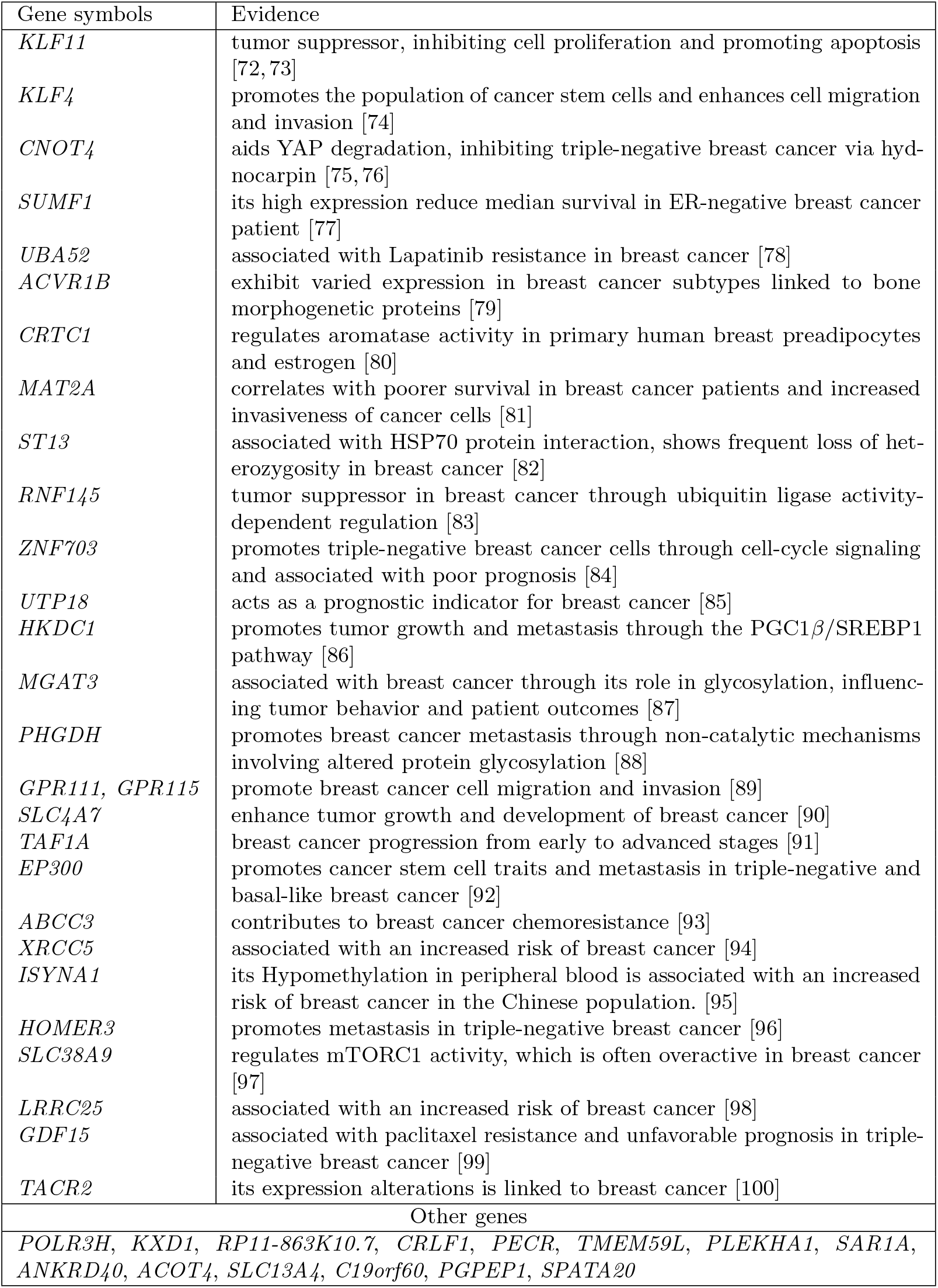
The potential breast cancer risk genes within 10kb of loci obtained through eQTL analysis, identified by MuGLasso or/and SMuGLasso and not the adjusted GWAS, found to be associated with breast cancer risk or tumor growth in the literature.

**S13 Table.** DTF3 loci detected by SMuGLasso and MuGLasso on *Arabidopsis thaliana* dataset. We present the genes identified through physical mapping of SNPs selected as associated with flowering time in *Arabidopsis thaliana* using SMuGLasso, MuGLasso and Adjusted GWAS.

| Methods | List of selected genes |
| --- | --- |
| Adjusted GWAS | **AT5G10100, AT5G45830, AT4G00730, AT4G00752, AT4G00630, AT4G00740, AT4G00750.** |
| SMuGLasso | **AT5G10100, AT5G45830, AT4G00730, AT4G00752, AT4G00630, AT4G00740, AT4G00750,** AT4G01915, AT5G15020, AT5G17710, AT5G27945, AT5G53410, AT3G58590, AT1G20130, AT3G29450, AT3G14490, AT1G03365, AT3G14470, AT1G28410, AT2G23430, AT3G27040, AT4G17970, AT4G09160, AT2G34890, AT4G30100, AT2G39990, AT4G35080, AT2G18500, AT3G46340, AT1G29300, AT3G27670, AT5G41820, AT2G38720, AT3G44610, AT4G33760, AT5G40450, AT1G27520, AT3G26140, AT4G16990, AT1G61360, AT3G61170, AT5G55910, AT2G25940, AT5G51830, AT1G43600, AT2G39310, AT4G34310, AT1G78970. |
| MuGLasso | **AT5G10100, AT5G45830, AT4G00730, AT4G00752, AT4G00630, AT4G00740, AT4G00750,** AT4G01915, AT5G15020, AT5G17710, AT5G27945, AT5G53410, AT3G58590, AT1G20130, AT3G29450, AT3G14490, AT1G03365, AT3G14470, AT1G28410, AT2G23430, AT3G27040, AT4G17970, AT4G09160, AT2G34890, AT4G30100, AT2G39990, AT4G35080, AT2G18500, AT3G46340, AT1G29300, AT3G27670, AT5G41820, AT2G38720, AT3G44610, AT4G33760, AT5G40450, AT1G27520, AT3G26140, AT4G16990, AT1G61360, AT1G43600, AT2G39310, AT4G34310, AT1G78970, AT4G33480, AT5G40290, AT1G12970, AT3G13550, AT2G32170, AT4G27290, AT1G59690, AT3G61170 AT5G55910, AT2G25940, AT5G51830. |

**S14 Table.**
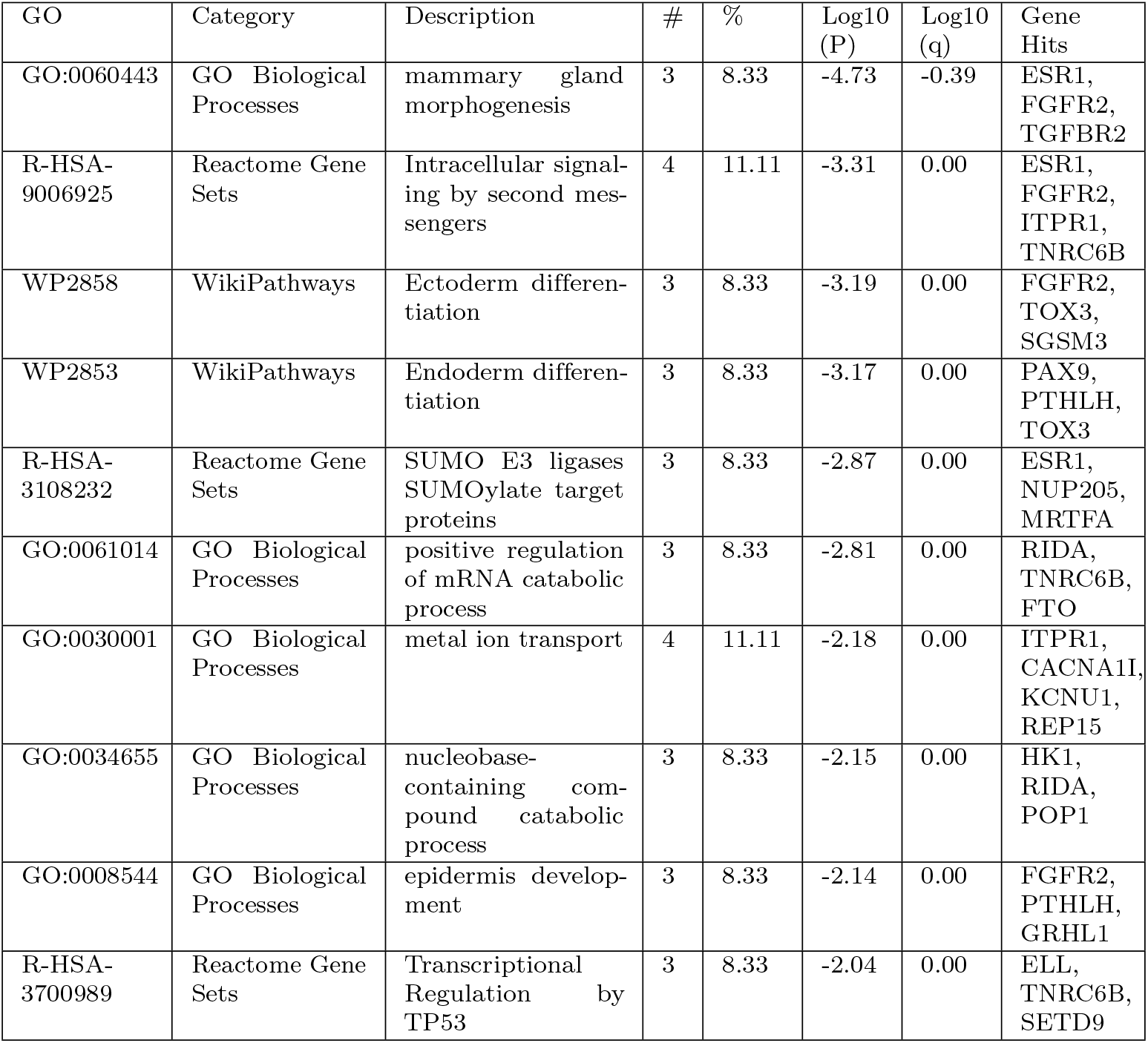
Summary of pathway and process enrichment analysis: Top 10 clusters of enriched terms, each described by one representative enriched term. “#” is the number of genes in the user-provided lists with membership in the given ontology term. “%” is the percentage of genes selected by SMuGLasso that are found in the given ontology term (only input genes with at least one ontology term annotation are included in the calculation). “Log10(P)” is the p-value in log base 10. “Log10(q)” is the multi-test adjusted p-value in log base 10.

**S15 Table.**
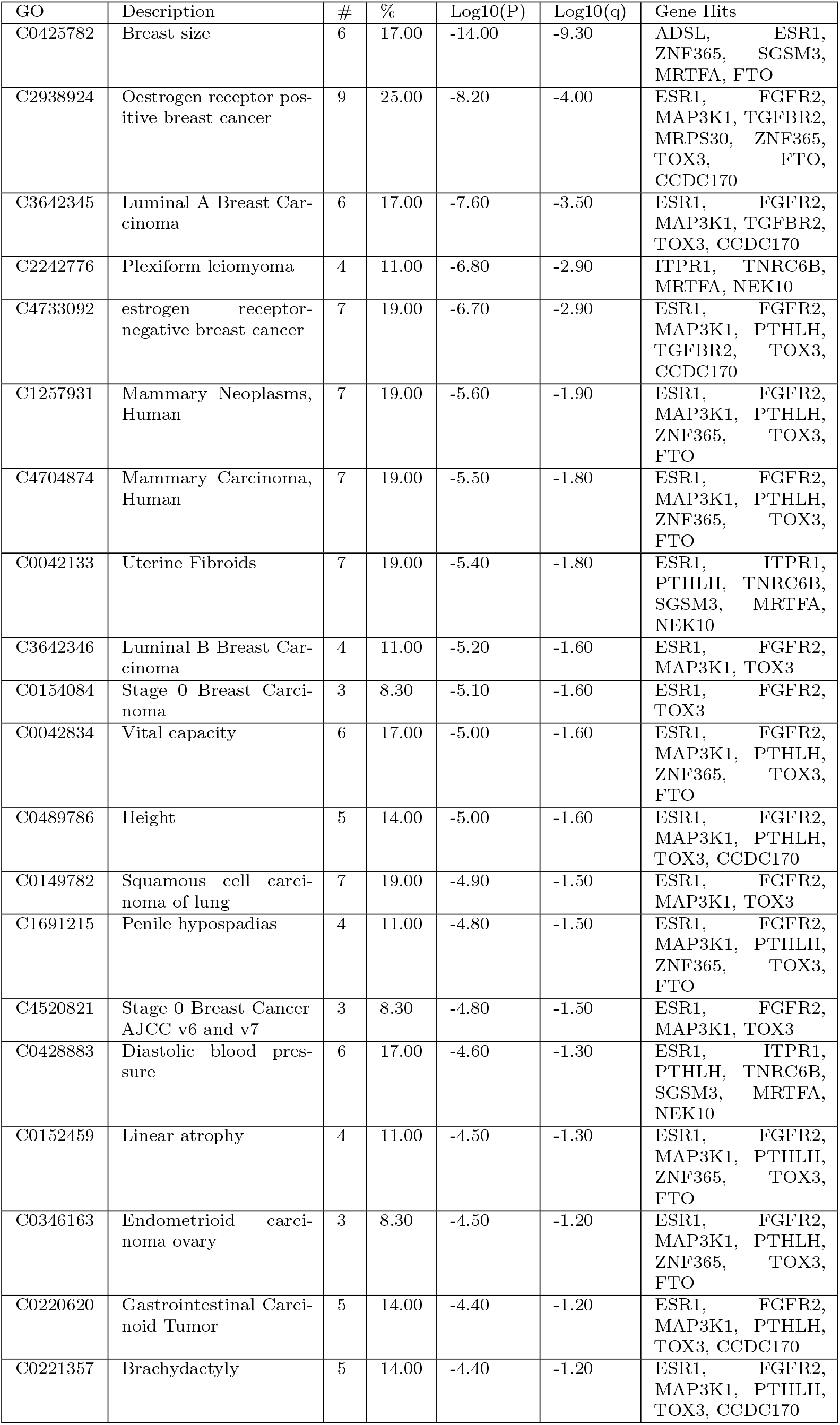
Summary of enrichment analysis in DisGeNET

**S16 Table.** Summary of enrichment analysis in Cell Type Signatures

| GO | Description | # | % | Log10(P) | Log10(q) | Gene Hits |
| --- | --- | --- | --- | --- | --- | --- |
| M40312 | DESCARTES FETAL THYMUS THYMIC EPITHELIAL CELLS | 5 | 14.00 | -5.00 | -1.60 | FGFR2, PAX9, CACNA1I, ASTN2, TOX3 |
| M39241 | LAKE ADULT KIDNEY C22 ENDOTHELIAL CELLS GLOMERULAR CAPILLARIES | 3 | 8.30 | -3.20 | -0.41 | EBF1, TGFBR2, PPFIBP1 |
| M39026 | FAN EMBRYONIC CTX EX 4 EXCITATORY NEURON | 3 | 8.30 | -3.00 | -0.31 | ITPR1, PPFIBP1, CACNA1I |
| M39234 | LAKE ADULT KIDNEY C15 CONNECTING TUBULE | 3 | 8.30 | -3.00 | -0.31 | ITPR1, PPFIBP1, TOX3 |
| M40237 | DESCARTES FETAL LUNG SQUAMOUS EPITHELIAL CELLS | 3 | 8.30 | -2.70 | -0.16 | PAX9, ZNF365, GRHL1 |
| M39233 | LAKE ADULT KIDNEY C14 DISTAL CONVOLUTED TUBULE | 3 | 8.30 | -2.70 | -0.16 | ITPR1, TNRC6B, TOX3 |
| M39136 | GAO ESOPHAGUS 25W C1 CILIATED EPITHELIAL CELLS | 4 | 11.00 | -2.70 | -0.15 | ASTN2, CCDC170, NEK10, SSBP4 |
| M39222 | LAKE ADULT KIDNEY C3 PROXIMAL TUBULE EPITHELIAL CELLS S1 S2 | 3 | 8.30 | -2.60 | -0.12 | PPFIBP1, TNRC6B, ASTN2 |

**S17 Table.**
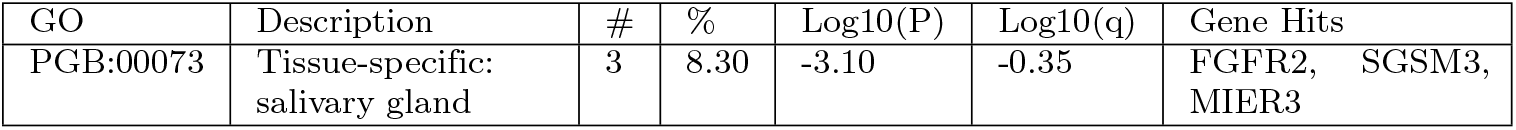
Summary of enrichment analysis in PaGenBase

| GO | Description | # | % | Log10(P) | Log10(q) | Gene Hits |
| --- | --- | --- | --- | --- | --- | --- |
| PGB:00073 | Tissue-specific: salivary gland | 3 | 8.30 | -3.10 | -0.35 | FGFR2, SGSM3, MIER3 |

**S18 Table.**
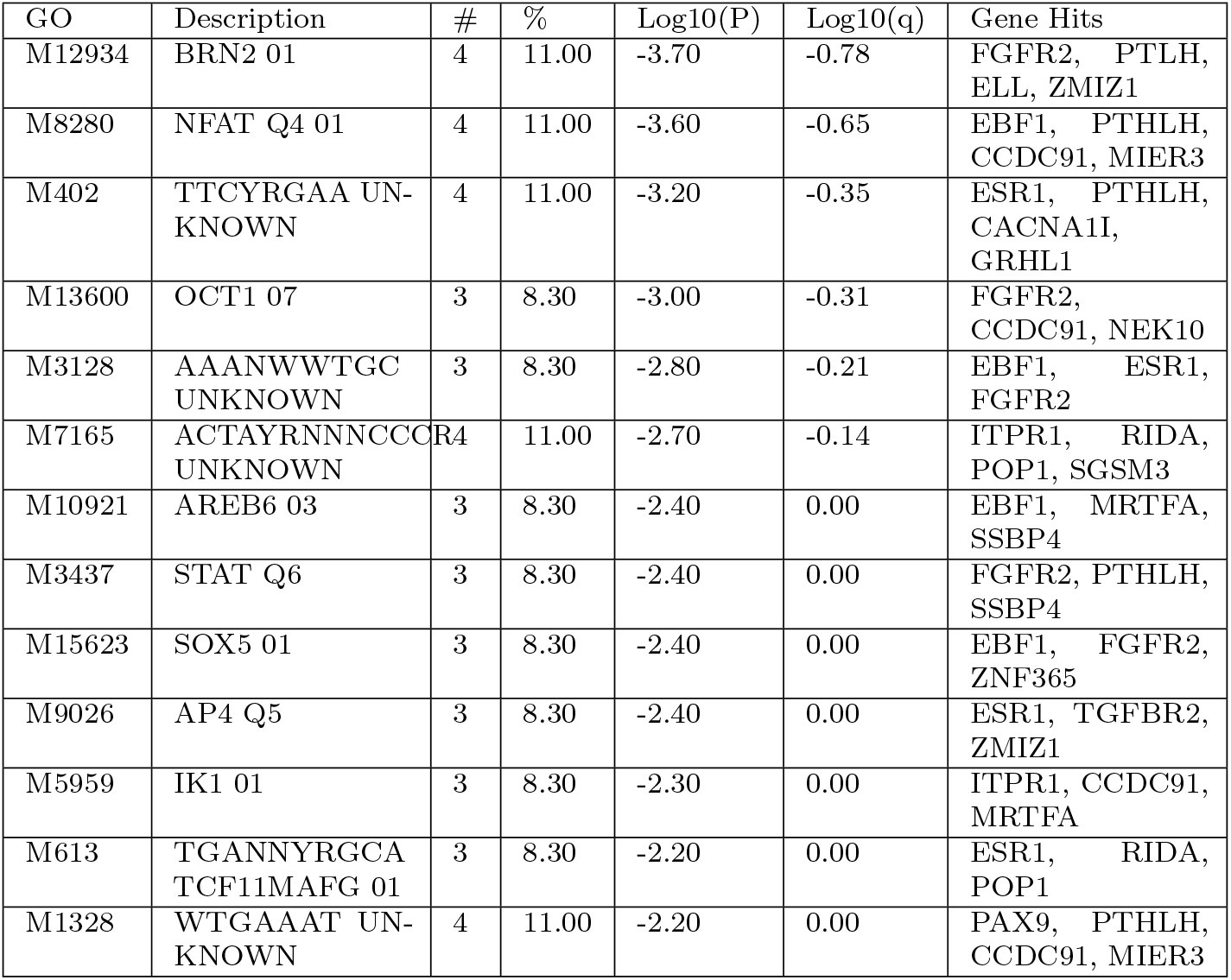
Summary of enrichment analysis in Transcription Factor Targets

**S19 Fig.**
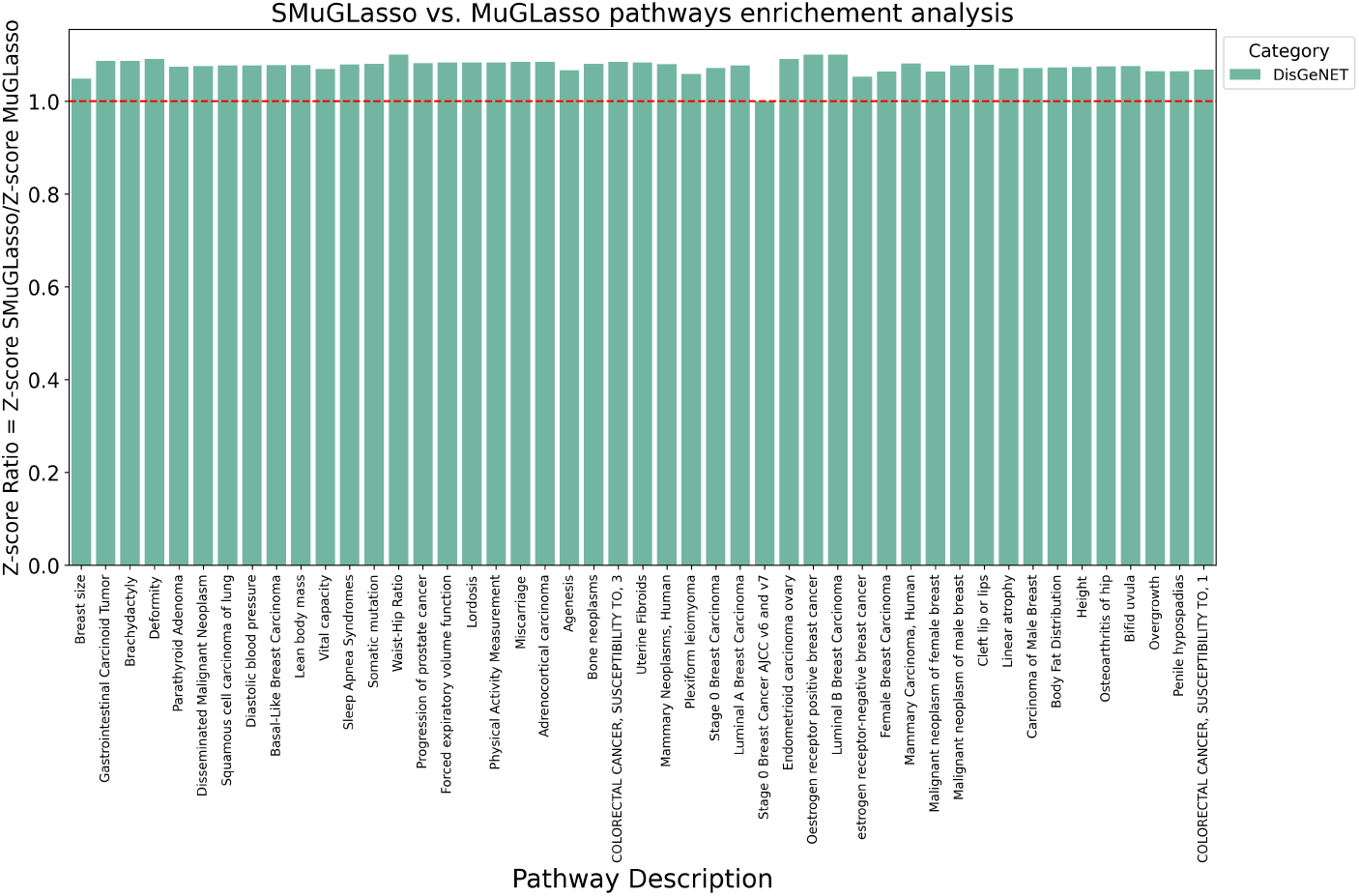
On DRIVE, Comparison of DisGeNET gene set enrichment between SMuGLasso and MuGLasso based on Z-score ratios. Bar heights represent the ratio of Z-scores (SMuGLasso/MuGLasso) for top DisGeNET common gene sets.

**S20 Fig.**
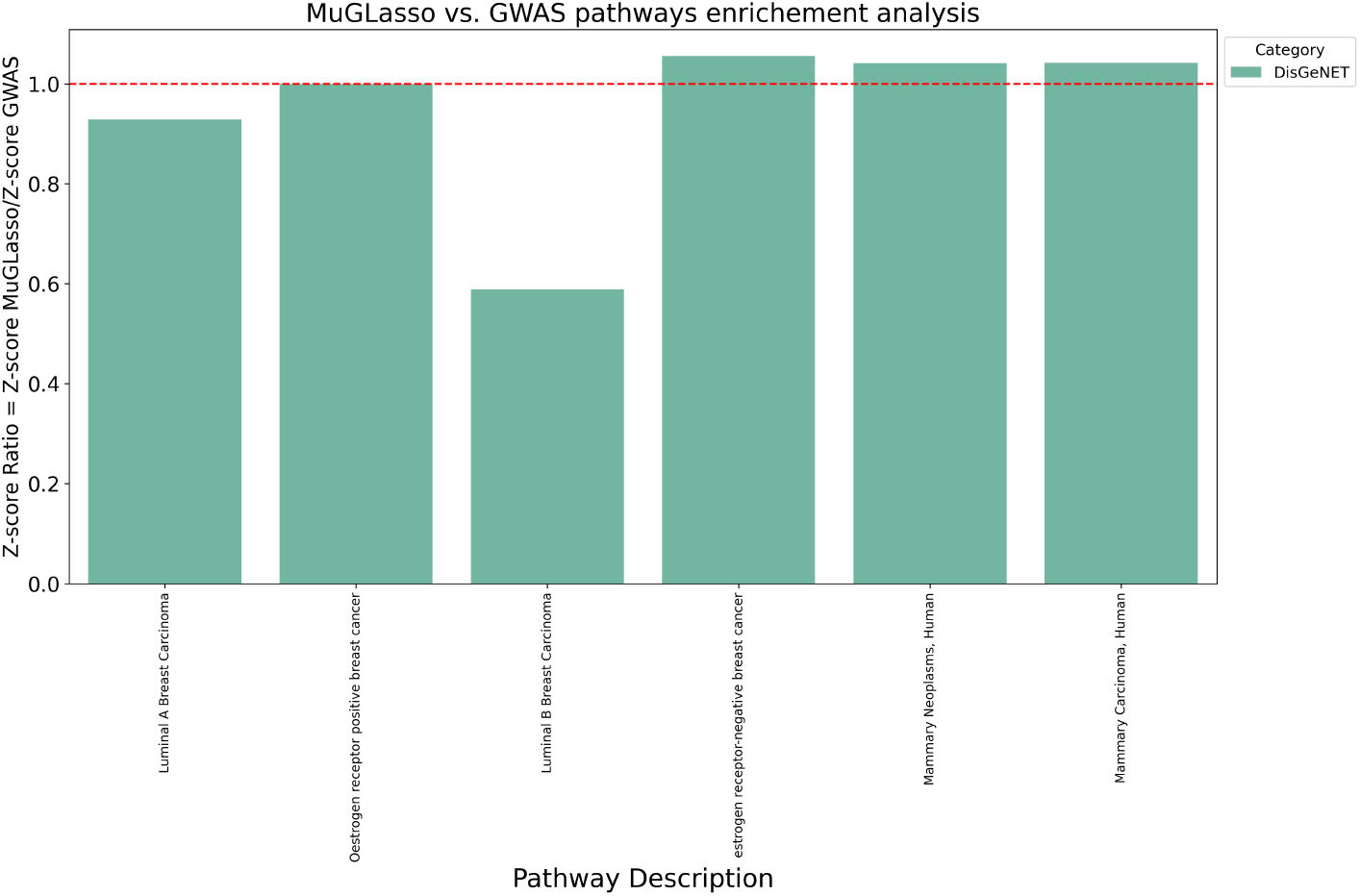
On DRIVE, Comparison of DisGeNET gene sets enrichment between MuGLasso and Adjusted GWAS based on Z-score ratios. Bar heights represent the ratio of Z-scores (MuGLasso/Adjusted GWAS) for top DisGeNET common gene sets.

**S21 Fig.**
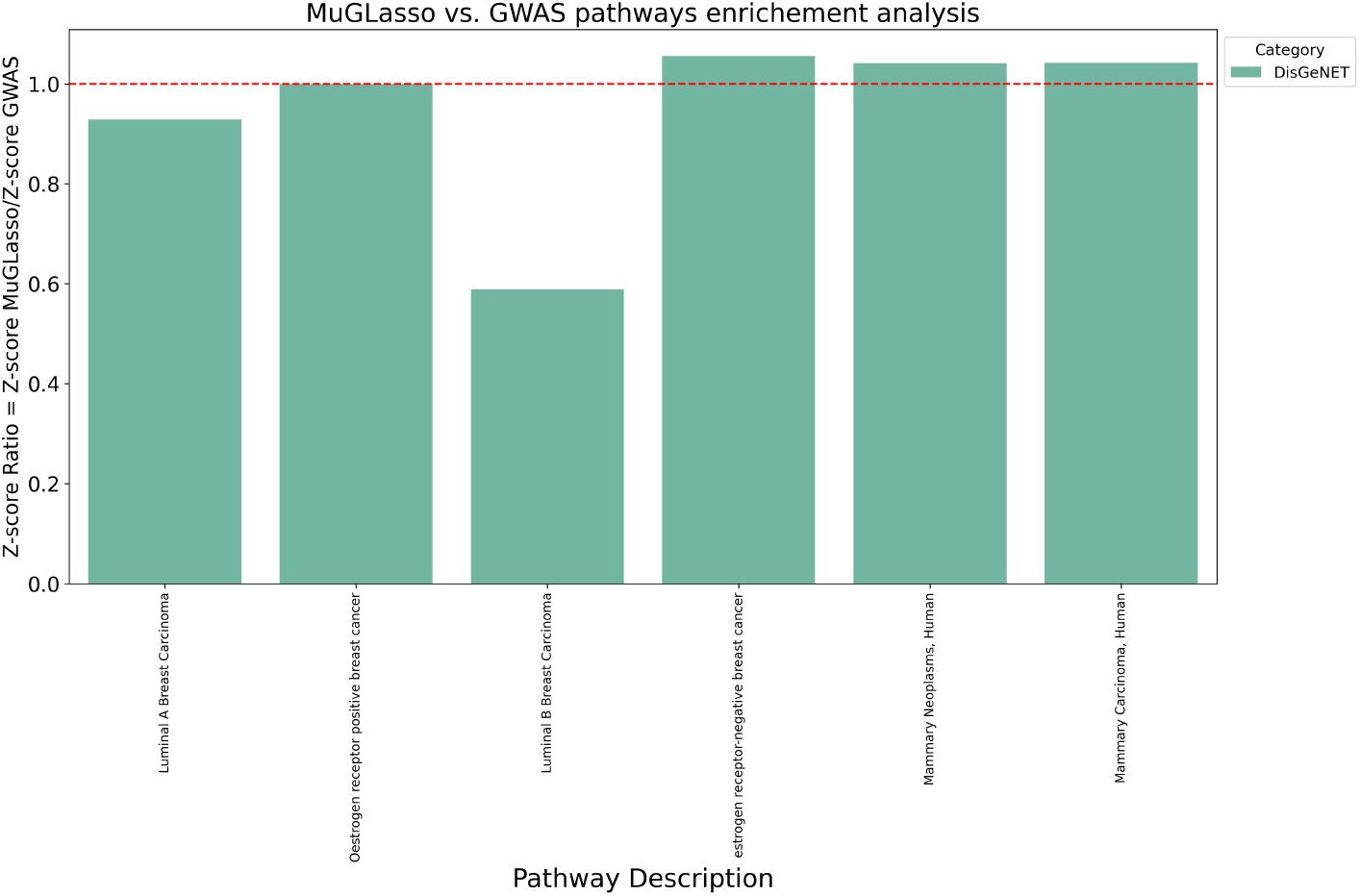
On DRIVE, Comparison of DisGeNET gene sets enrichment between MuGLasso and Adjusted GWAS based on Z-score ratios. Bar heights represent the ratio of Z-scores (MuGLasso/Adjusted GWAS) for top DisGeNET common gene sets.

**S22 Fig.**
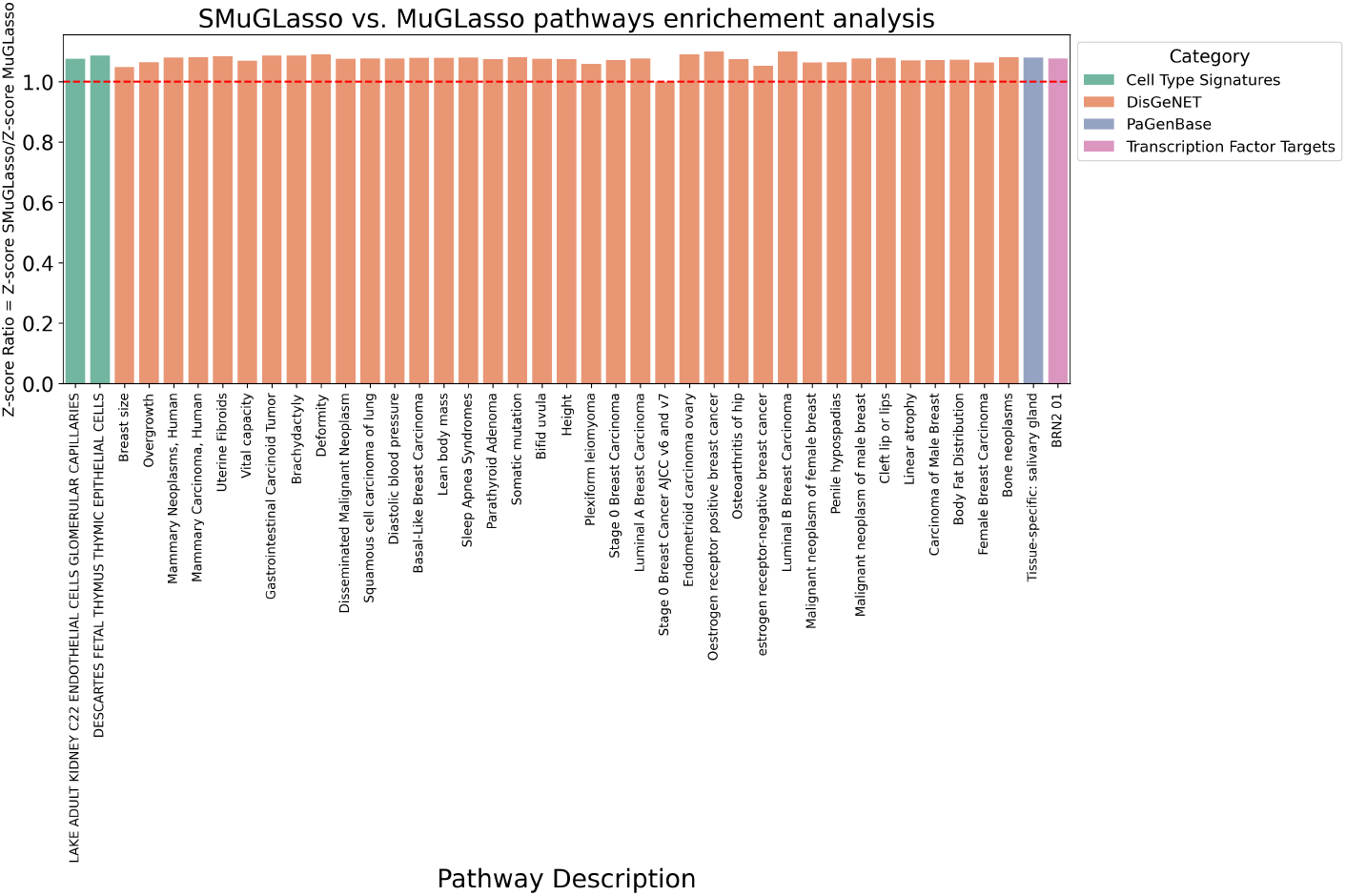
On DRIVE, Comparison of pathway enrichment between SMuGLasso and MuGLasso based on Z-score ratios. Bar heights represent the ratio of Z-scores (SMuGLasso/MuGLasso) for top DisGenet common gene sets.

**S23 Fig.**
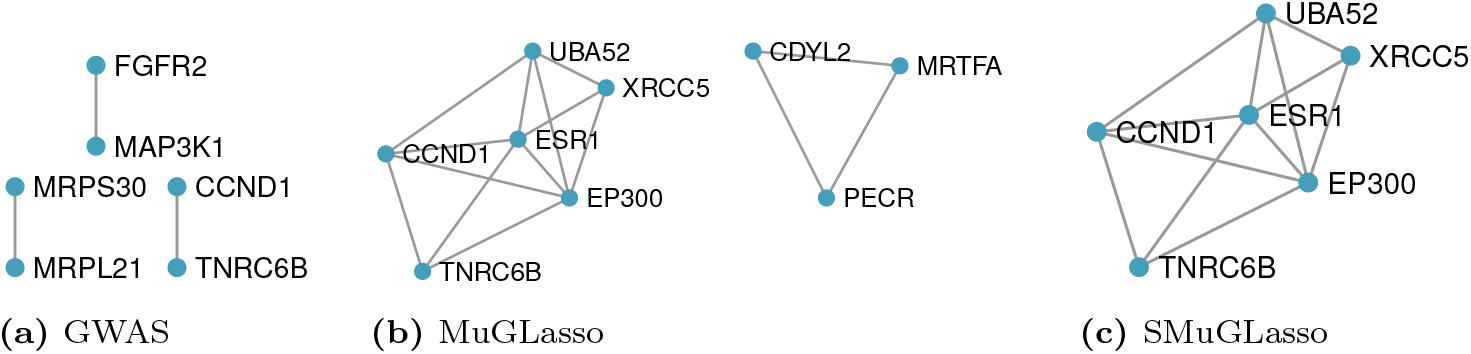
Modules of the PPI of known interactions between genes identified through physical and eQTL mapping of the SNPs selected by Adjusted GWAS, SMuGLasso and MuGLasso on DRIVE.

## Acknowledgments

The authors would like to thank Adeline Fermanian, Vivien Goepp, Héctor Climente-González, Gwenaëlle Lemoine, Antoine Poirier and Lotfi Slim for fruitful discussion. This work was supported by the French Agence Nationale de la Recherche (ANR-18-CE45-0021-01 and ANR-19-P3IA-0001). OncoArray genotyping and phenotype data harmonization for the Discovery, Biology, and Risk of Inherited Variants in Breast Cancer (DRIVE) breast-cancer case control samples was supported by X01 HG007491 and U19 CA148065 and by Cancer Research UK (C1287/A16563).

